# The biochemical composition of coronavirus Nucleocapsid proteins drives the bicistronic properties of their sub-genomic mRNAs

**DOI:** 10.64898/2026.11.04.764307

**Authors:** Federico De Angelis, Hou Wei Chook, Ternenge T. Apaa, Amanda H. Seekings, Lorraine McElhinney, Marike Geldenhuys, Rochelle Rademan, Wanda Markotter, Rute M. Pinto, Lu Lu, Eleanor Gaunt, Samantha J. Lycett, Paul Digard

**Affiliations:** The Roslin Institute, The University of Edinburgh, Easter Bush, Midlothian EH25 9RG, UK; Rabies & Viral Zoonoses, Animal and Plant Health Agency, Addlestone, Surrey, KT15 3NB; Department of Animal Science, School of Agriculture, Policy and Development, University of Reading, Whiteknights, Reading RG6 6EU, UK; Centre for Viral Zoonoses, Department of Medical Virology, Faculty of Health Sciences, University of Pretoria, South Africa

## Abstract

The nucleocapsid (N) gene of coronaviruses (CoVs) encodes a structural RNA-binding protein required for viral assembly. In SARS- and MERS-CoV viruses, reading frame 2 of N contains an overlapping open reading frame (ORF) encoding an innate immune antagonist accessory protein. However, the extent of this dual coding capacity across the CoV family, and the genetic features driving overlapping ORF formation, remain unexamined. Here, we developed an *in-silico* tool (ORFan) to screen structural gene sub-genomic mRNA sequences from 152 species of coronavirus for the presence of overlapping accessory gene candidates. We found that frame 2 of the N gene in Alpha-, Beta- and Delta-CoVs frequently contains a potential accessory ORF (AcORF), conserved at the subgenus level, unlike the spike, envelope and membrane genes. We also show that CoV nucleotide biases in conjunction with codon biases integral to N protein function drive the relatively high abundances of start codons in frame two and stop codons in frame three, explaining the tendency towards frame 2 AcORFs. We further suggest that viral nucleotide biases and the codon composition required for RNA-binding function facilitate overlapping ORF formation in other virus families.

## INTRODUCTION

Coronaviruses (CoVs) are a diverse family of single stranded, positive sense RNA viruses that infect birds and mammals. They represent health and financial threats due to the wide range of diseases they cause and their spillover potential, as emphasized by the global crises caused by the coronavirus disease 2019 (COVID-19) pandemic and the severe acute respiratory syndrome (SARS) and Middle East respiratory syndrome (MERS) epidemics^1,2,3,4,5,6,7^.

CoVs are divided into four genera, *Alphacoronavirus* (α-CoVs), *Betacoronavirus* (β-CoVs), *Gammacoronavirus* (γ-CoVs), and *Deltacoronavirus* (δ-CoVs), each further divided into subgenera based on phylogenetic relatedness^8^. All CoV genomes have the same basic 5’-3’ structure: Leader, 5’ untranslated region (5’UTR), replicase and virulence factors (ORF1a/1ab), spike (S), envelope (E), membrane (M), nucleocapsid (N) and a 3’UTR with a poly adenosine (poly-A) tail^9^. Interspersed between and sometimes contained within the essential structural genes S, E, M and N, smaller, less well-conserved accessory open reading frames (AcORFs) which encode additional virulence factors can be found^10,11,12^. While the polyproteins ORF1a and 1ab are synthesised by ribosomes that directly translate the CoV genome upon cell entry, downstream genes encoding the structural and accessory proteins are translated from individual sub-genomic messenger RNA (sgmRNA) molecules^13^. SgmRNAs are synthesised via discontinuous transcription, a form of RNA recombination directed by repeating transcription regulatory sequences (TRSs) located in the 5′ UTR (the leader TRS-L) and upstream of most ORFs that follow ORF1ab (in the genome ‘body’ - TRS-Bs)^14,15^. As the viral RNA polymerase complex synthesises the antisense strand of the genome, it may stall at a TRS-B, allowing the nascent antisense RNA molecule to detach from its template and reanneal with the complementary TRS-L for transcription re-initiation. The leader sequence in the 3′ of the subgenomic RNAs then primes positive strand sgmRNA synthesis, producing a nested set of sgmRNAs sharing identical 5’ and 3’ ends.

In CoVs, the N gene is the last of the structural genes, proximal to the 3’ end of the viral genome^16^. It expresses the nucleocapsid protein, a structural protein that binds viral RNA, forming a ribonucleoprotein (RNP) complex, necessary for correct genome packaging during virion assembly^17,18^. The N protein has a modular organisation, with two evolutionarily conserved functional domains spaced between three less-conserved intrinsically disordered regions (IDRs)^19^. The conserved domains are an N-terminal RNA-binding domain (RBD) and a C-terminal dimerization domain (DD), required to promote RNP polymerisation^17^. The IDRs are located at the N- and C-termini of the protein, and in between the functional domains following a serine/arginine-rich flexible linker (LKR) region that contains several phosphorylation sites^20,21^.

The N sgmRNA of viruses such as SARS-CoV and SARS-CoV-2^22,23^, MERS-CoV^24,25,26^, bovine CoV (BCoV)^27^, murine hepatitis virus (MHV)^28^ and porcine deltacoronavirus (PDCoV)^29,30^ also contain AcORFs embedded in the +1 reading frame (frame 2) of the N gene which are surmised to be expressed via ribosomal leaky scanning. This is the phenomenon in which an initiation codon (usually methionine) in an mRNA molecule is skipped by scanning ribosomes during translation initiation, leading to initiation at a downstream initiation site, also typically an AUG^31^. The efficiency of this process is affected by multiple factors, including the "Kozak" consensus sequences flanking the canonical and downstream initiation sites ^32,33,34,35^, as well as 5’UTR length^36,37^.

The gene products of the currently identified AcORFs nested within N have evidence of involvement in host-pathogen interactions^38,22,24,27,28,30^. For instance, ORF8b in MERS-CoV has been suggested to suppress the interferon-β induction cascade by competing with IKKε for binding to the 70-kDa heat shock protein (HSP70)^24^. NS7 from PDCoV has also been found to target IKKε to hinder its interaction with both TRAF3 and IRF3^29^. ORF9b from SARS-CoV localises to mitochondria and targets MAVS, affecting the MAVS/TRAF3/TRAF6 signalosome^38^ while ORF9b from SARS-CoV-2 has been described to localise to the mitochondrial external membrane and interact with MAVS by associating with translocase of the mitochondrial outer membrane 70 (Tom70) protein^23^.

Thus, there is a growing body of evidence from clinically and/or economically important CoVs indicating that the CoV N gene can be functionally bi-cistronic, but a comprehensive survey of N-encoded AcORFs is lacking, and the partial annotation of most CoV genomes makes it difficult to obtain a full picture of the overall distribution and conservation of AcORFs embedded in the essential genes. In this study, we performed a systematic search for AcORFs in CoV structural genes across 152 viral species by developing a novel bioinformatic tool (‘ORFan’) parameterised to assess leaky ribosomal scanning potential. We found that in general, CoV N genes contain many more candidate AcORFs than the S, M and E genes, driven by N gene nucleotide composition and a lower content of codons encoding hydrophobic amino acids.

## RESULTS

To perform a systematic search for AcORFs in alternative reading frames of CoV N genes, we first developed a database of full-length coronavirus genome sequences available from NCBI Virus, supplemented with further unpublished sequences from bat CoV surveillance programmes in the UK and South Africa. The final dataset contained 152 coronavirus species: 55 α-CoVs, 74 β-CoVs, 15 δ-CoVs and nine γ-CoVs (**Supplementary Fig 1a, Supplementary Table 2**). We included sequences to represent the diversity of the *Orthocoronavirinae* subfamily, although the uneven distribution of viruses across the four genera and 25 subgenera reflects the number of publicly available distinct CoV species for which the full sequence could be found at the time of the analysis. Eighty-nine out of the 152 CoV sequences were of bat origin (**Supplementary Fig 1b),** consistent with the role of the *Chiroptera* order as a CoV reservoir^39,40^. Other host groupings with multiple CoV species included artiodactyls (*n* = 13), primates (*n* = 8), rodents (*n* = 7) and birds (*Anseriformes n* = 6; *Passeriformes n* = 6).

To investigate the coding capacity of N, we manually identified the TRS-L, the N gene TRS (TRS-B-N) and gene body sequences, and assembled them to form N-specific sub-genomic mRNA (sgmRNA) sequences for each virus in our dataset (**Fig 1a)**. TRS-L and B-N sequences for each subgenus (**Supplementary Table 3**) were identified based on previous publications^41,42^. To identify potential AcORFs embedded in the N gene, we developed an <u>O</u>pen <u>R</u>eading <u>F</u>rame <u>an</u>alyser (ORFan) bioinformatic tool, which scores ORFs in alternative reading frames of a given gene based on the likelihood of a translation initiation event occurring at their start codon. ORFan works in two steps: firstly, it annotates all ORFs in an mRNA sequence that start with an AUG codon and terminate with a stop codon; then secondly, after defining the longest ORF as the primary gene, the likelihood of translation initiation at another ORF occurring to produce a functional polypeptide (the latter defined by a minimum ORF length) is calculated (**Fig 1b**). Briefly, for a given ORF, ORFan defines its length in nucleotides, the Kozak^31,32^ context of its start codon, the number of intervening start codons between the ORF and the 5’UTR of the main gene, their Kozak contexts and the ORF distance from the 5’UTR. Each of these factors are assigned a numerical value (see **Methods**) and used to calculate an overall score that reflects the likelihood of the ORF to be accessed by ribosomes via leaky ribosomal scanning. An ORFan score ≥0.1 suggests a good accessory gene candidate, 0.1>x≥0.01 as moderate, 0.01>x≥0.001 as poor and <0.001 indicates no candidate. To test ORFan sensitivity, it was applied to other instances of positive sense, negative sense and dsRNA virus genes that are known to have a 2^nd^ reading frame AcORF embedded in another gene thought to be expressed via leaky ribosomal scanning (**Supplementary Table 1**). To test specificity, the sequence sets included strains with documented non-functional AcORFs. ORFan identified the majority of these AcORFs as good candidates, while mostly failing to do so in sequences previously annotated as lacking them (**Supplementary Fig 2a**). The exception was two Tilapia Lake virus sequences previously annotated as not possessing an intact S9.3 gene sequence^108^. However, one has a C-terminally truncated ORF which still met the minimum length threshold, and while the other lacks the normal opening AUG codon, ORFan identified the next one downstream as still accessible to leaky ribosomal scanning.

**Figure 1.**
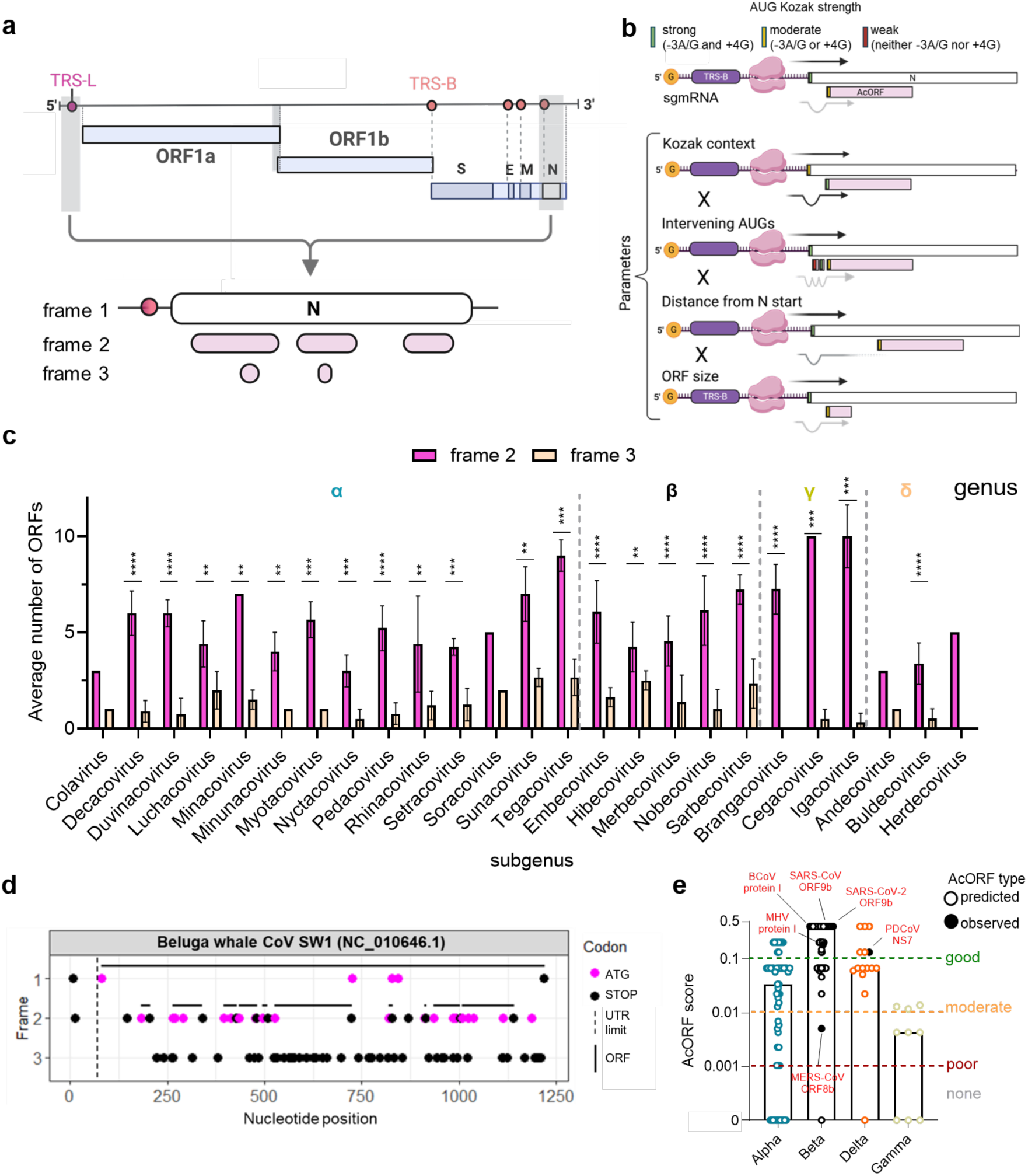
Frame 2 of the N gene of α-, β- and δ-CoVs encodes ORFs likely to be expressed as accessory genes. a) *In silico* pipeline for the generation of N sub-genomic mRNA sequences and the identification of all ORFs in the three positive reading frames. b) Diagrammatic summary of parameters included in ORFan AcORF likelihood calculations. c) Average number of ORFs identified in the second or third reading frame of the N gene of CoV subgenera. Error bars represent the standard deviation. Averages of frame 2 and 3 were compared by paired t-test (*p*-values: *<0.05; **<0.01; ***<0.001; ****<0.0001). d) Distribution of ATG and Stop codons within Beluga whale CoV SW1 N sgmRNA across reading frames 1-3. Solid lines represent ORFs, the dashed line delimits the 5’UTR. e) Median AcORF likelihood score divided by CoV genus. An ORFan score ≥ 0.1 suggests a good (green line) accessory gene candidate. 0.1>x≥0.01 suggests a moderate (yellow line), 0.01>x≥0.001 suggests a poor (red line) and <0.001 was considered as no candidate. Predicted AcORF candidates are shown in open circles, AcORFs with known protein products are shown in black and annotated on the figure.

Applying ORFan to our dataset of CoV sequences showed that frame 2 of the N gene contained between two and 10 times more ORFs than frame 3, depending on CoV subgenus (**Fig 1c).** To visualise potential AcORFs in the CoV N genes, the ORFan outputs were used to draw ORF coordinate maps for all N sgmRNAs included in this study. As expected, frame 1 of all viruses contained one long ORF for the N gene (**Fig 1d; Supplementary Fig 3**). In contrast, frame 3 was often rich in stop codons and poor in start codons, leading to a few small or even no ORFs. Frame 2 tended to have fewer stop codons than frame 3 (although more than frame 1) and many more start codons, giving a variety of ORF structures. In total, ORFan found 1015 ORFs (810 – frame 2; 205 - frame 3) in the N gene alternative reading frames. One hundred and thirty-four of those were classified as potential accessory genes (ORFan score > 0.001), and all of them resided in N frame 2. β-CoVs had the greatest number of good AcORF candidates (ORFan score ≥ 0.1), followed by δ-CoVs and α-CoVs (**Fig 1e**). These results were corroborated by the fact that ORFan successfully identified all six previously observed N AcORFs as *bona fide* genes, suggesting that N gene bicistronicity could be a common feature of the α-, β- and δ-CoV families.

We then compared ORFan’s performance against a previously published method for identifying accessory genes in alternative reading frames^43^. Applying the Frameshift (FS) pipeline to our N gene dataset identified 44 ORFs with a *p*-value ≤ 0.05, against the 134 identified by ORFan. The 44 statistically significant AcORF candidates included three previously characterised N AcORFs, against the six identified by ORFan (**Supplementary Fig 2b**). FS did not classify any sarbecovirus ORF9bs or MERS-CoV ORF8b as statistically significant, despite evidence for their expression in infected cells^44,45,46,25,47,48^. Conversely, all AcORF candidates identified by FS were also identified by ORFan, suggesting ORFan has higher sensitivity for accessory gene prediction, especially for shorter candidates.

### N AcORF score distribution is linked to viral genetic relatedness

To look for associations between CoV N gene phylogenetic relatedness and their respective AcORF score, a maximum likelihood phylogenetic tree was generated using a multiple sequence alignment of all CoV N gene sequences. The phylogeny showed clear segregations of species into groups (**Fig 2)** corresponding to the International Committee on Taxonomy of Viruses subgenus classification of coronaviruses^8^. The analysis also showed that the distribution of good, moderate or poor N accessory gene candidates tended to cluster at the subgenus level. In β-CoV subgenera for example, all embecoviruses, most nobecoviruses and 75% of sarbecoviruses had a good predicted AcORF candidate. Merbecoviruses had a majority (80%) of moderate AcORF candidates and three out of four hibecoviruses had a good AcORF candidate. The δ-CoVs mostly had moderate AcORF candidates, while α-CoV subgenera showed the highest degree of intra-subgenus variability. Nevertheless, trends of AcORF conservation (or lack thereof) could be observed in six α-CoV subgenera (*Pedacovirus*, *Luchacovirus*, *Rhinacovirus*, *Myotacovirus*, *Sunacovirus*, and *Setracovirus*), in which most viruses had AcORF candidates with similar likelihoods of expression. Host distribution analyses showed that four out of five β-CoV subgenera have a largely monophyletic host group, as well as seven of the 14 α-CoV ones. Some subgenera, (*e.g.* β-CoV *Embecovirus*, α-CoV *Luchacovirus*, *Duvinacovirus* and *Tegacovirus* and δ-CoV *Buldecovirus)* had a broader host-range, spanning three or more host genera. No clear correlation was seen between host type and AcORF candidate score. Overall, these results suggest that the distribution of AcORF candidates is associated with CoV phylogeny but not host evolutionary group.

**Figure 2.**
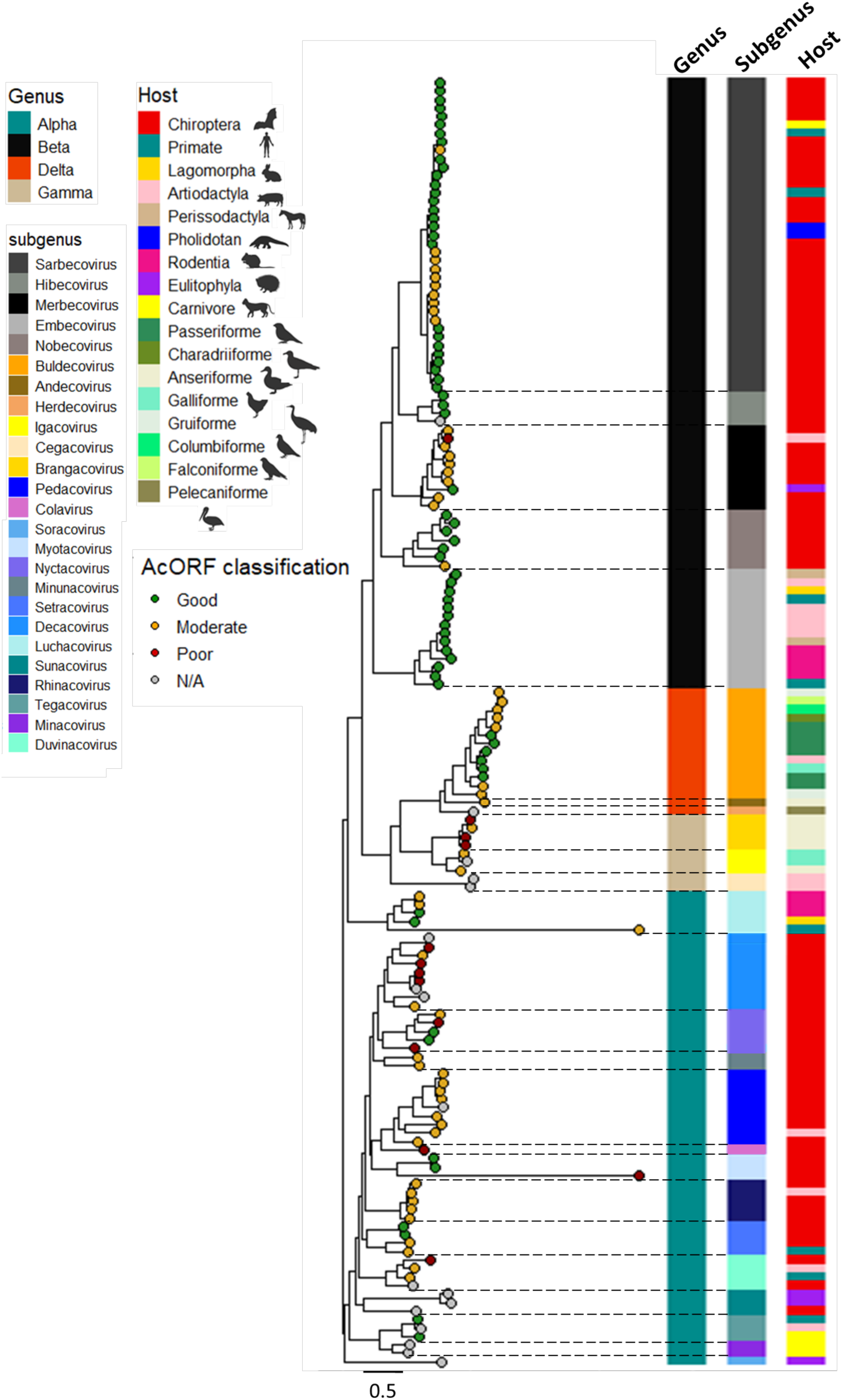
N AcORF score distribution is linked to viral genetic relatedness. Midpoint-rooted maximum likelihood tree showing the phylogenetic relationships of 152 coronavirus N genes. Tree scale represents the number of nucleotide substitutions. Tips are colour-coded based on the likelihood of encoding a good, moderate, poor or no accessory gene candidate in frame 2 of N. Bar colours indicate genus, subgenus and host annotation. The phylogenetic tree was generated on IQTree using the GTR+I+G model with 1500 ultrafast bootstrap replicates.

### AcORFs are often conserved among subgenera, but not at the family level

Structurally, N has two conserved functional domains: RNA binding (RBD) and dimerization (DD) domains (**Supplementary Fig 4a**). These are flanked by intrinsically disordered regions (IDRs) and interspaced by a serine-arginine-rich linker region and another IDR^16,17,49^. To determine whether AcORF conservation could be tied to conservation of the N functional domains, a CoV family-wide N protein conservation profile was generated by aligning all N sequences in our dataset and extracting their amino acid biochemical conservation scores^50^. At the pan-CoV level, only 17 amino acids were entirely conserved, and all of them resided in the functional domains (**Supplementary Fig 4b**). We therefore turned to subgenus-level N alignments to probe the relationships between AcORF and N sequences.

Examining β-CoVs first, as the genus with the highest likelihood of encoding an AcORF; N conservation analyses showed a much higher degree of similarity at the subgenus level, with the highest conservation observed in the functional domains and the IDRs being generally more variable (**Fig 3a-e**). All members of the β-CoV subgenera (except for MERS-CoV and one nobecovirus) possessed AcORFs with start codons in the N-terminal IDR, which despite IDR variability, mostly showed striking conservation of both position and Kozak consensus. The position of AcORF stop codons was more variable, although mostly conserved within subgenera as to whether they lay within the N RBD, (sarbecoviruses and nobecoviruses) or in the central SR/IDR domain (merbecoviruses, embecoviruses and hibecoviruses). The corresponding analyses of AcORF structure for the other genera of CoVs generally showed more variation at the subgenus level, perhaps reflecting fewer individual sequences, greater overall diversity and overall lower probabilities of containing an AcORF (**Supplementary Figure 5**), although α-CoV luchacoviruses, nyctacoviruses and rhinacoviruses, δ-CoV buldecoviruses and y-CoV brangacoviruses subgenera showed some consistency.

**Figure 3.**
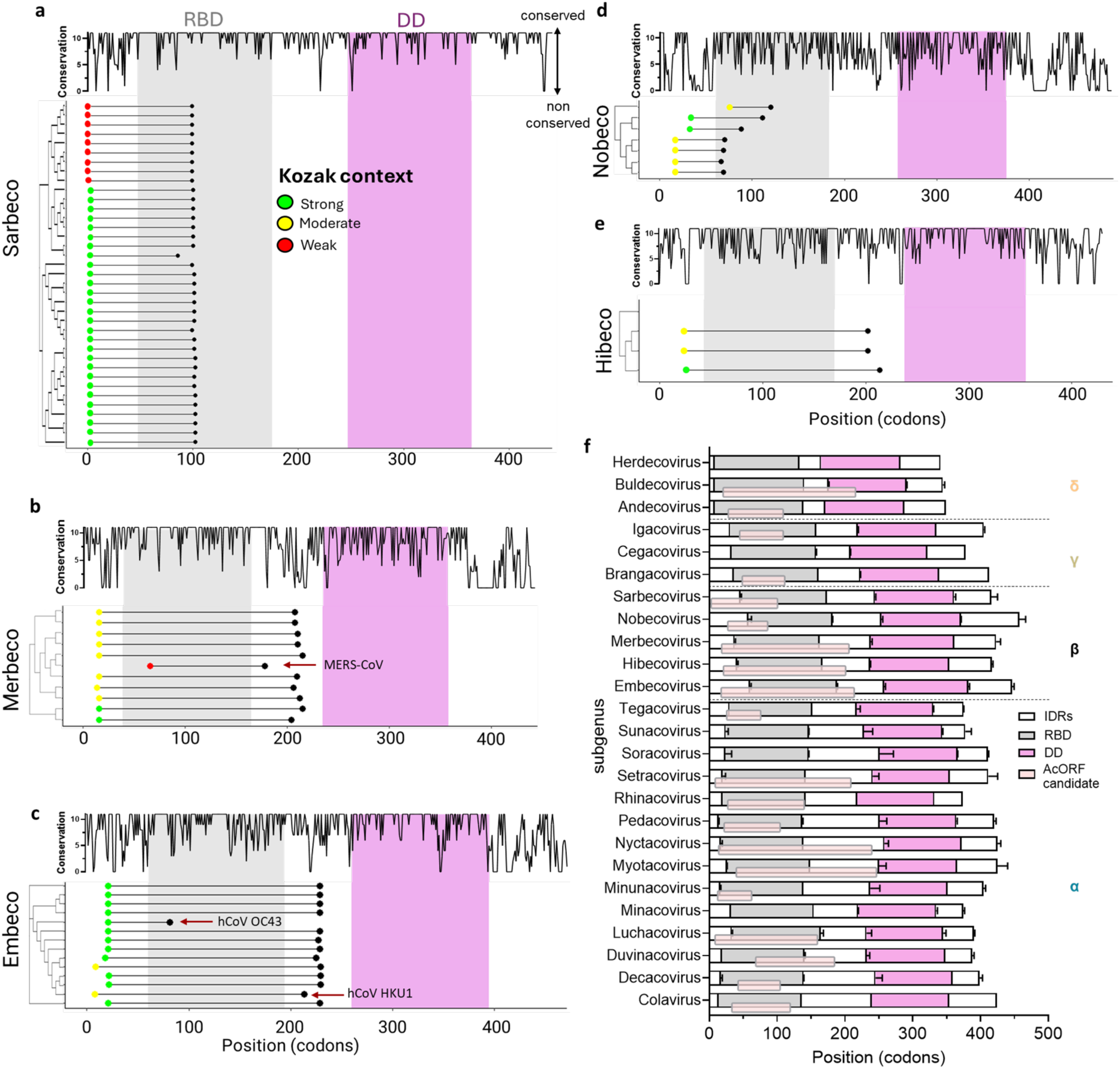
AcORF coordinate analysis suggests conservation at the subgenus level. a) *Sarbecovirus*, b) *Merbecoviru*s, c) *Embecovirus*, d) *Nobecovirus* and e) *Hibecovirus* N protein residue-by-residue biochemical conservation profile and species-specific frame 2 AcORF codon coordinates. The median coordinates of N RBD (grey) and DD (fuchsia) regions for each sub-genus and the codon coordinates of individual AcORFs were plotted using RStudio. Cladograms were generated using MEGA11. Start codons of AcORFs and N genes are coloured according to their Kozak context (green = strong; yellow = moderate; red = weak). f) Conservation of N functional domain and AcORF coordinates. Average codon coordinates of N structural modules and AcORF candidates (pink boxes) within each subgenus are plotted. Error bars represent the standard deviation.

To visualise the overall pattern of AcORF conservation in relation to N across all CoV subgenera, the coordinates of the functional domains of N were extracted and averaged between each subgenus to generate subgenus-specific N functional maps (**Figure 3f**), and the average subgenus-specific AcORF candidates codon coordinate mapped onto them. This showed a diversity of IDR lengths across the CoV family, but that as expected for a model in which a second cistron is accessed by ribosomal leaky scanning, AcORF candidates consistently overlapped the N RBD in all subgenera. Beyond that, there was no clear pattern of whether AcORFs started within the N-terminal IDR or N, or ended in N or the internal IDR, or even the N DD coding region. Thus, although the coordinates of N-encoded AcORFs are often conserved at the subgenus level, their gene starts and stops are not correlated with a particular domain of N.

### The distribution of frame 2 and 3 ORFs in CoV N genes is due to nucleotide and codon biases in N

To determine the genetic factors that drive the elevated number of ORFs in frame 2 of CoV N genes compared to frame 3, the nucleotide distributions at position one, two and three of each codon within reading frame 1 of the N genes in our dataset were calculated. Results showed that, on average, CoV N genes overrepresent G (31%) and A (30%) at position one (**Fig 4a**), A (34%) and C (27%) at position two (**Fig 4b**) and U (36%) at position three (**Fig 4c**), independently of genus. Codon frequencies for frames 1, 2 and 3 were also calculated, showing that none of the ten most common codons in CoV Ns (which amount to ∼34% of the whole gene) possess a U at position two (a necessary requirement for the formation of a stop codon in frame 2), while the fourth (A<u>AU</u> - 3.57%) and seventh (G<u>AU</u> – 3.23%) most frequent codons could lead to the formation of a frame 2 start codon (**Fig 4d**). Indeed, the codons CUG (4.45%), AUG (3.96%) and UUG (3.58%) were the three most abundant codons in N frame 2, while every stop codon was found to be below the 1.56% random distribution threshold (UGA = 1.11%; UAG = 0.68%; UAA = 0.63%). On the other hand, stop codons <u>U</u>GA (4.75%) and <u>U</u>AA (3.9%) were the first and third most frequent codon in frame three, respectively. Overall, these results suggest that the relative lack of N-U-N codons in N frame 1, as well as the elevated frequency of AAU and GAU codons contribute to the high number of putative ORFs in frame 2 and their simultaneous suppression in frame 3.

**Figure 4.**
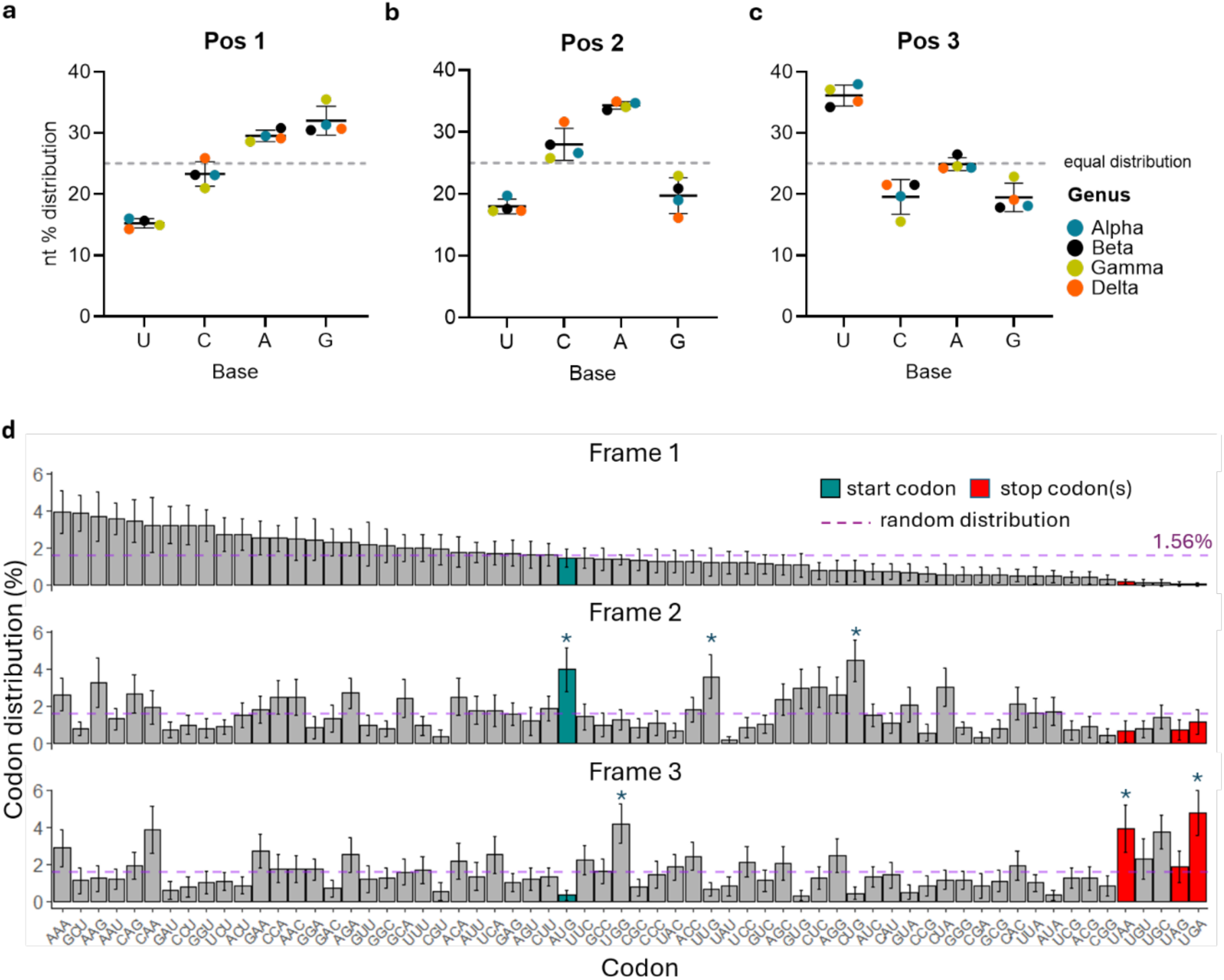
The observed ORF distribution in N reading frames 2 and 3 is caused by nucleotide biases in N. a) Average nucleotide frequency by frame 1 (N) codons at position one, b) two and c) three, coloured by genus. Error bars represent the SEM. The dotted line indicates the expected frequency given equal nucleotide composition. d) Mean frequency of codons in N frames 1, 2 and 3, ranked according to frame 1. Error bars represent the standard deviation. Start and stop codons are highlighted in cyan and red respectively. The three most frequent codons in frame 2 and 3 are highlighted by asterisks. The random distribution threshold (1/64 = 1.56%) is represented by dashed purple lines.

### N genes are more likely to contain potential AcORFs than other CoV structural genes due to a lower hydrophobic amino acid content

To determine whether the dual coding capacity of N is common to all CoV structural genes, we extracted the S, E and M genes from each complete virus genome (137 in total; **Supplementary Table 2**) in our dataset, compiled sgmRNA sequences as before (**Supplementary Table 3**) and repeated the ORFan analyses. This showed the same bias against ORFs in frame 3 as in the N gene, but also that the S gene had by far the highest number of ORFs in alternative reading frames among the CoV structural genes, including N (**Fig 5a; Supplementary Table 4**). N had the second highest number, followed by M and then E, consistent with the overall size of each gene. However, the AcORF-to-ORF ratio as assessed by ORFan showed that on average, an individual CoV N gene (ratio value: 0.14) is ∼15 times more likely to contain an AcORF than S (0.009), ∼150 times more than E (0.0009), and ∼three times more than M (0.054) (**Fig 5b**). CoV genomes have been described to overrepresent Us while suppressing Cs and Gs^51,52,74,96^. Overall, the individual S, E and M genes follow this pattern, but N has a significantly lower U content, balanced by slightly higher amounts of A, C and G (**Fig 5c**). Lower U content in N was particularly marked at frame 1 positions 1 and 2, especially in comparison to the E and M genes (**Supplementary Fig 6).** Differences in nucleotide distribution could not be attributed to large differences in relative synonymous codon usage (RCSU), as the four structural genes had comparable values for most U-containing codons, except that N had the lowest frequency of N-U-A codons, which are required for the formation of two of the three stop codons in frame 2 (**Supplementary Fig 7**). We therefore analysed the average amino acid composition of the four CoV structural proteins, which showed that N contains a significantly lower percentage of hydrophobic amino acids and a higher content of polar, acidic and basic amino acids (**Fig 5d, Supplementary Fig 8**). Except for tyrosine (Y), tryptophan (W) and alanine (A), hydrophobic amino acids tend to be encoded by codons with U at position two and are therefore more likely to cause a stop codon (UAA, UGA, UAG) in frame 2 compared to other codons (**Fig 5e**). To have a frame 2 AcORF accessible through leaky ribosomal scanning, structural genes need to therefore avoid codons encoding hydrophobic amino acids in their N-termini to prevent formation of stop codons, while favouring N-A-U codons which enable the formation of start codons, such as CAU (histidine – H), GAU (aspartic acid – D), AAU (asparagine – N) and UAU (Y). We therefore aligned all S, E, M and N protein sequences in our dataset and looked at the amino acid distribution within the first 200 positions (approximately the region occupied by N AcORFs) of each alignment. For E (between 75-109 amino acids long), we aligned the whole sequence. In S, the first 50 amino acids showed a high frequency of hydrophobic amino acids mostly interspersed by either polar or special case amino acids **(Fig 5f)**. E and M had a high frequency of hydrophobic amino acids along the whole length of their alignments (**Fig 5g, h**). In contrast, the first 150 positions in the N alignment were predominantly occupied by polar, basic and acidic amino acids, except for two pairs of two consecutive hydrophobic amino acids at position 95-96 and 132-133 (**Fig 5i**). Overall, these results suggest that the lower content of hydrophobic amino acids in the N-terminal domains of N, combined with a high content of polar, basic and acidic amino acids drive the birth and retention of AcORFs in frame 2 of the gene.

**Figure 5.**
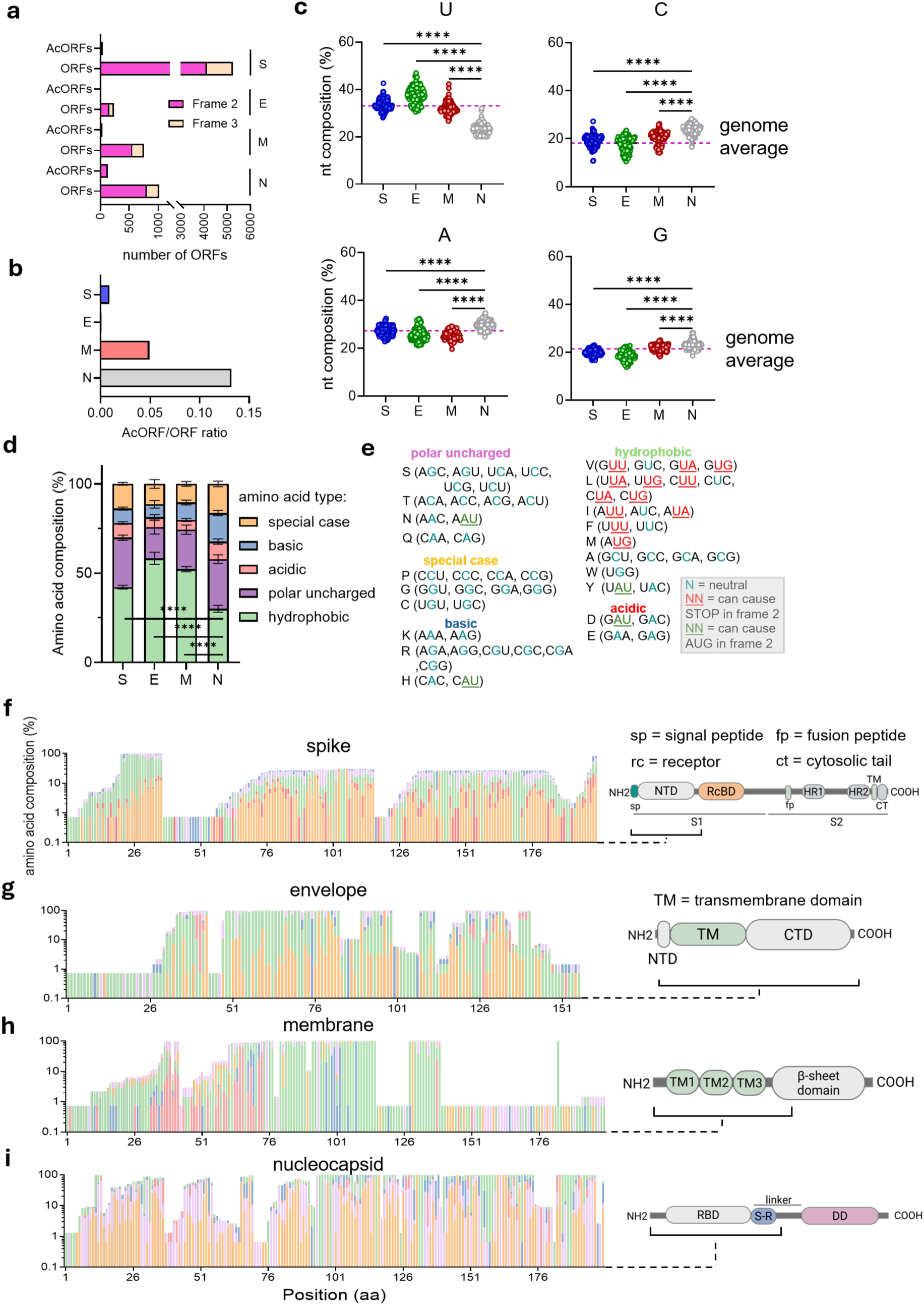
N is more likely to encode accessory genes than other CoV structural genes due to lower hydrophobic amino acid content. a) Number of ORFs vs AcORFs identified in frame 2 and 3 of CoV structural genes. b) AcORF encoding probability of S, E M and N calculated as AcORF-to-ORF ratios. c) Nucleotide frequency percentage of S, E, M and N genes in our dataset. Dashed lines represent the average nucleotide frequency across all CoV genomes. Statistical significance was calculated via One-Way ANOVA with Bonferroni correction for multiple comparisons (ns: non-significant, *: p<0.05; **: p<0.01; ***: p<0.001; ****: p<0.0001). d) Amino acid distribution of S, E, M and N grouped by biochemical properties. e) Amino acid codons, grouped by biochemical properties, annotated based on their likelihood of producing a start codon (green), stop codon (red) or anything else (neutral; blue) in frame 2. (f-i) Amino acid type distribution by position in protein multiple sequence alignments for (f) Spike, (g) Envelope, (h) Membrane and (i) Nucleocapsid. Diagrams on the right indicate protein domain structure and the regions plotted on the left. Note that the y axis does not always sum to 100% because of variations in protein length.

## DISCUSSION

Several CoVs, including SARS-CoV^53,54^, SARS-CoV-2^55,56^, MERS-CoV^24,25,26^, PDCoV^29,30^ and MHV^28,38^ are known to contain AcORFs in the second reading frame of their N genes. Although there are examples where an internal TRS-B within N produces a separate sgmRNA^15,30^ the majority of these characterised N AcORFs are expressed or surmised to be expressed via leaky ribosomal scanning^46,57^. Here, we used data from studies on translation initiation^33,34,58^, ribosomal scanning^36,59^, and minimal viral ORF lengths^60,61,62^ to develop ORFan, an *in-silico* tool that identifies embedded ORFs and scores their likelihood of being expressed via ribosomal leaky scanning, and then used it to systematically screen a library of CoV structural genes for evidence of an AcORF. We found that N has the highest density of ORFs in its second reading frame, while being virtually devoid of them in its third reading frame due to a scarcity of start codons and an overabundance of stop codons. ORFan scores showed that α-, β- and δ-CoVs often possess N genes containing ORFs with a moderate to high likelihood of being expressed during infection in their second reading frame, while γ-CoVs had the lowest mean likelihood score, despite having the highest average number of frame 2 ORFs. A possible reason for the difference between γ-CoVs and other genera is that the N gene of all γ-CoVs analysed is in an optimal Kozak context, therefore decreasing the probability of downstream AUG initiation via leaky ribosomal scanning.

ORFan appeared to be more sensitive than the Frameshift (FS) pipeline developed by Schlub et al.^43^, managing to identify all seven described N-encoded AcORFs, against the three identified by FS. However, while FS scores are skewed towards longer ORFs, the tool is agnostic towards expression mechanism, while ORFan is biased towards leaky ribosomal scanning, and does not consider other translation initiation mechanisms. Tandem use of these tools can therefore be an effective way to look for new AcORF candidates in other viral families. Of note, both FS and ORFan identified clusters of AcORF candidates in α- and β-CoVs that have not been described before, strengthening the idea that N-encoded AcORFs are a family-wide characteristic of CoVs. Importantly, ORFan has a minimum ORF size cut-off point of 120 nucleotides (39 amino acids plus a stop codon), which is within the range of the smallest known proteins expressed by mammalian viruses^60,61,62,63,64^, but ribo-seq data on other viruses^65,66^ and recent efforts on human cells^67^ suggest that ORFs as small as 10 amino acids can express a functional peptide, indicating the ORFan size cut-off is conservative.

Phylogenetic analyses revealed that closely related CoVs tend to encode AcORFs with similar likelihoods of expression, suggesting that genetic relatedness is a more relevant factor than host type for the presence or absence of AcORFs in N. These observations agree with the accepted notion that some CoV AcORFs are conserved at the subgenus level^12,68^ but also represent the first evidence of *Merbecovirus*, *Embecovirus* and *Buldecovirus*-wide N-encoded AcORF conservation, as well as the first evidence for N-encoded AcORFs in α-CoVs.

Spatial distribution analyses of AcORFs in the different β-CoV subgenera showed that although most AcORFs partially or entirely overlap N RBDs, their start and stop codons often dwell outside of the RBD boundaries and are highly conserved at subgenus level. Given the higher degree of conservation of the N RBD compared to the flanking IDRs, these results seem counterintuitive. Nevertheless, they agree with previous research suggesting that overlapping regions in dual-coding RNA virus genes often encode for highly disordered proteins or protein regions^53,69,70^. They are also consistent with selection pressure to maintain the AcORFs.

Lastly, nucleotide and protein composition analyses of CoVs structural genes showed that N is a better frame 2 AcORF reservoir than S, E and M due to its specific nucleotide and amino acid biases. Our findings agree with previous work which showed that viral overlapping and non-overlapping genes differ in nucleotide and amino acid composition but disagree with the suggestion that overlapping genes are depleted in A nucleotides, as well as ApU and UpA dinucleotides^71^. N has a higher A content than S, E and M, and its 1^st^ and 3^rd^ most abundant frame 1 codons are AAA (K) and AAG (K). However, different families of RNA viruses exhibit varying forms of genome composition bias^72,73,74,75^. CoVs overrepresent codons ending in U or A across their genome^76^. This remains true for N, as highlighted by our RSCU analysis. The higher content of adenosines in N may be important for additional reasons; in some contexts, adenosines are selected for as they enable ADAR-based editing, which replaces mRNA-encoded adenosines with inosine, mimicking host cell transcripts and subverting dsRNA-mediated immune sensing^97,98,99,100^. At the protein level, the abundant usage of K residues in the N-terminal region of N implies that its functional role may be a contributing factor in influencing AcORF formation in alternative reading frames. The high content of basic (K, R, H) and polar uncharged (Q, N, T, S) amino acids in N are important for its overall function as an RNA binding protein (which requires positive charges)^17,19,21,77,78,79^ as well as its cytosolic localisation (whereas S, E and M travel through vesicular compartments)^80,81,82^. Together, these nucleotide- and codon-level constraints create a distinctive genetic landscape within the coronavirus N gene that is unusually permissive to the emergence and maintenance of AcORFs.

These insights further enhance our understanding of the coding capacity of coronaviruses, but more broadly suggest that there might be AcORF reservoir ‘candidate’ genes in other viruses, based on cognate protein function and overall genome coding biases. Other viral families may follow a similar trend to what we observe here. Bunyaviruses^83,84,85^, rotaviruses^86,87^, influenza A^88,89,90^ and astroviruses^91^ possess bi-cistronic genes in which the main ORF encodes a protein with RNA-binding properties, suggesting that nucleotide-binding function could be a favourable factor in the evolution of overlapping ORFs.

Additionally, the coding biases that regulate AcORF ‘birth’ and retention also impact AcORF protein composition. The second reading frame of N has a high prevalence of codons encoding hydrophobic amino acids. Approximately one third of all sequence-encoded across the tree of life are membrane proteins^109^. Most transmembrane proteins contain α-helical or β-barrel domains that cross the membrane lipid bilayer^101^, while shorter amphipathic helices can insert into one leaflet of the membrane and associate with it through their hydrophobic surface^110^. ORF9b from SARS-CoV-2 and NS7 from PDCoV have been described to localise to mitochondria^23, 104^, which are membrane-bound organelles, and similar phenotypes can be observed in non-coronavirus accessory proteins too. PB1-F2 from Influenza A and NSP6 from species A rotavirus localise to the mitochondria^105,106^, while human astrovirus ORFx acts as a viroporin^107^. Based on these observations, we would argue that viral protein function, as well as genome-wide nucleotide biases, should be important factors to incorporate in the identification of bicistronic genes in other virus families. Lastly, we advise that further efforts should focus on determining the expression and functional role of the AcORF candidates identified in this study, as they will contribute to a broader understanding of CoVs infectious cycles.

## MATERIALS AND METHODS

### Datasets generation

Twelve segment 2 IAV sequences, 11 bunyavirus S segments, 12 segment 11 rotavirus A sequences and 10 segment 9 Tilapia lake virus sequences were downloaded from NCBI virus (**Supplementary Table 1**). For astroviruses, the whole genomes of 11 astrovirus species were downloaded and the ORF2 sequences extracted in FASTA format (**Supplementary Table 1**).

Coronavirus full genome sequences were acquired from the National Centre for Biotechnology Information (NCBI) Virus (https://www.ncbi.nlm.nih.gov/labs/virus/vssi/) database (data acquired from September 2021 to September 2025) and supplemented with 14 unpublished N gene sequences arising from bat coronavirus surveillance programmes in the UK and South Africa. Structural gene nucleotide sequences (S, E, M and N) were extracted in FASTA format and named according to the viral species they belonged to.

Additional sequence metadata is available in **Supplementary Table 2**.

### TRS identification

The viral genomes in our dataset were manually scanned to identify the TRS-L and TRS-B for the structural genes S, E, M and N (**Supplementary Table 3)**. The regions spanning the 5’ end of each viral genome and their respective TRS-L were then merged at their TRS-B complementary points and added upstream of the corresponding structural gene sequence to generate a representation of the sub-genomic mRNA that would be produced during viral replication.

### Alternative reading frame ORF mapping

For each CoV structural gene, concatenated FASTA files containing sgmRNA sequences were analysed with the R package “viralORF” (https://github.com/ChookHW/viralORF). An ORF is defined as any sequence starting with an AUG start codon and ending with an in-frame stop codon (UAA, UGA, UAG). The longest ORF in the positive-sense strand was extracted for further analysis.

### Scoring AcORF translation initiation potential with ORFan

For each ORF in frame 2 and 3 of S, E, M or N, the ORFan R script (**Supplementary Materials – ORFan script**) annotates Kozak strength, the number of intervening AUGs in the untranslated region between structural gene start and ORF start, the Kozak strength of each intervening start codon, the length of the ORF and the nucleotide distance between structural gene start and ORF start. The Kozak strengths of each start codon are turned into numerical values representing the *initiation potential* (*ip*) of each ORF based on the identity of the two nucleotides flanking the translation initiation site (TIS). *Ip* values were extracted from multiple studies that have experimentally determined how different Kozak sequences affect translational initiation *in vitro*^33,34,58^. *Ip* scores are assigned as follows: If two consensus nucleotides ( -3 A/G) and (+4 G) are present at the TIS, it is defined as a strong Kozak consensus, with an *ip* of 0.9. When either -3 (A/G) or +4 (G) is present it was considered a moderate initiation probability (score: 0.453333), and weak when neither -3 (A/G) nor +4 (G) is present (*ip* = 0.15). Conversely, the initiation probability of the N gene start, as well as any intervening AUG codons were turned into ‘skipping’ probability (*sp*; 1-*Ip* score) to estimate the chance of a ribosome missing the AUG and continuing to scan. ORFs are also assigned a *length (l)* score, to reflect the probability of their encoding a functional polypeptide. *L* values are assigned based on the size of small viral ORFs known to encode functional polypeptides, such as bovine papillomavirus E5 protein (45 codons)^60^, geminivirus ORF4 and ORF5 (54 and 46 codons respectively)^62^ or Kaposi’s sarcoma-associated herpesvirus vSP-1 (49 codons)^61^. Therefore, ORFs >=120 nucleotides (including the stop codon) are assigned an *l* score of 1, while ORFs < 120 nucleotides are scored as 0.

Additionally, ORFs are assigned a *distance* (*d*) score: If an ORF start is within 100 nucleotides from the structural gene start, then it is assigned a *d* score of 1; if the ORF start is found after 100 nucleotides but before 300 from the main gene start, then its *d* score is halved to 0.5; if the ORF is found >300 nucleotides after the main gene start site, its *d* score is 0. Distance scores are based on data suggesting that in human cells, longer 5’UTRs affect ribosome processivity and lead to a decrease in translational initiation^36^. AcORF scores are then calculated as: *AcORF translation likeli*ℎ*ood* = *ip x* (*sp*^1^ *x sp*^2^ *x sp^n^* … ) *x d x l*

Candidate AcORFs were then classified as good (score > 0.1); moderate (0.1 > score ≥ 0.01), poor (0.01 > score ≥ 0.001) or no candidate (score <0.001).

### ORFAn performance comparison

ORFan predictive precision was compared with another available bioinformatic pipeline that detects candidate overlapping AcORFs in viral genes from single sequences (the “frameshift” (FS) R package described in reference 43) (https://github.com/TimSchlub/Frameshift). All codes were run according to the authors’ instructions on RStudio v.2025.05.1+513 "Mariposa Orchid" for Windows.

### Phylogenetic analyses

N gene sequences were aligned using MUSCLE as a built-in function of the Molecular Evolutionary Genetics Analysis (MEGA11) software^92^. Phylogenetic trees were inferred via IQTree (http://www.iqtree.org/)^93^ using the maximum-likelihood method with the GTR+I+G substitution model and 1500 ultrafast bootstrap replicates. Trees were visualised and annotated using FigTree (v.1.4.4).

### Protein conservation analysis and domain mapping

N amino acid sequences were analysed and RNA-binding domain and dimerization domain coordinates extracted using InterPRO (https://www.ebi.ac.uk/interpro/)^94^ from the EMBL European Bioinformatics Institute. N amino acid sequences were aligned on Jalview (v.2.11.4.0)^95^ using MUltiple Sequence Comparison by Log-Expectation (MUSCLE) with the default settings. Sequence conservation scores at each amino acid position were extracted from Jalview and visualised using GraphPad Prism (v.10.2).

### Gene composition analyses

Codon distributions, relative synonymous codon usage (RSCU) values and nucleotide frequencies of the structural genes were extracted from codon aligned fasta files using MEGA11^92^.

### Statistical analyses

All statistical tests were performed in GraphPad Prism 10 Software. Specific statistical tests were chosen depending on the experimental design and are stated in each figure legend (ns: non-significant, *: p<0.05; **: p<0.01; ***: p<0.001; ****: p<0.0001).

## Supporting information

Supplementary information (Figures and smaller tables)

## Data availability

All underlying sequence data used in this study are publicly available via Genbank. The ORFan code and outputs are given in the Supplementary Information.

## Author Contributions

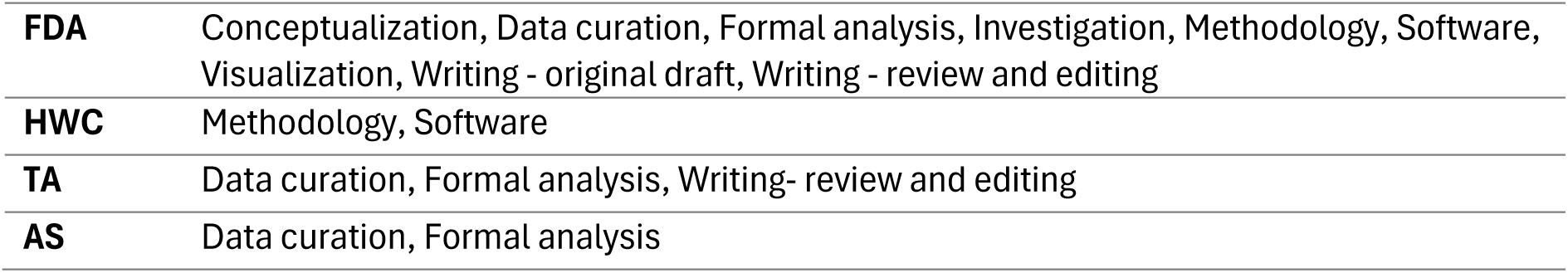
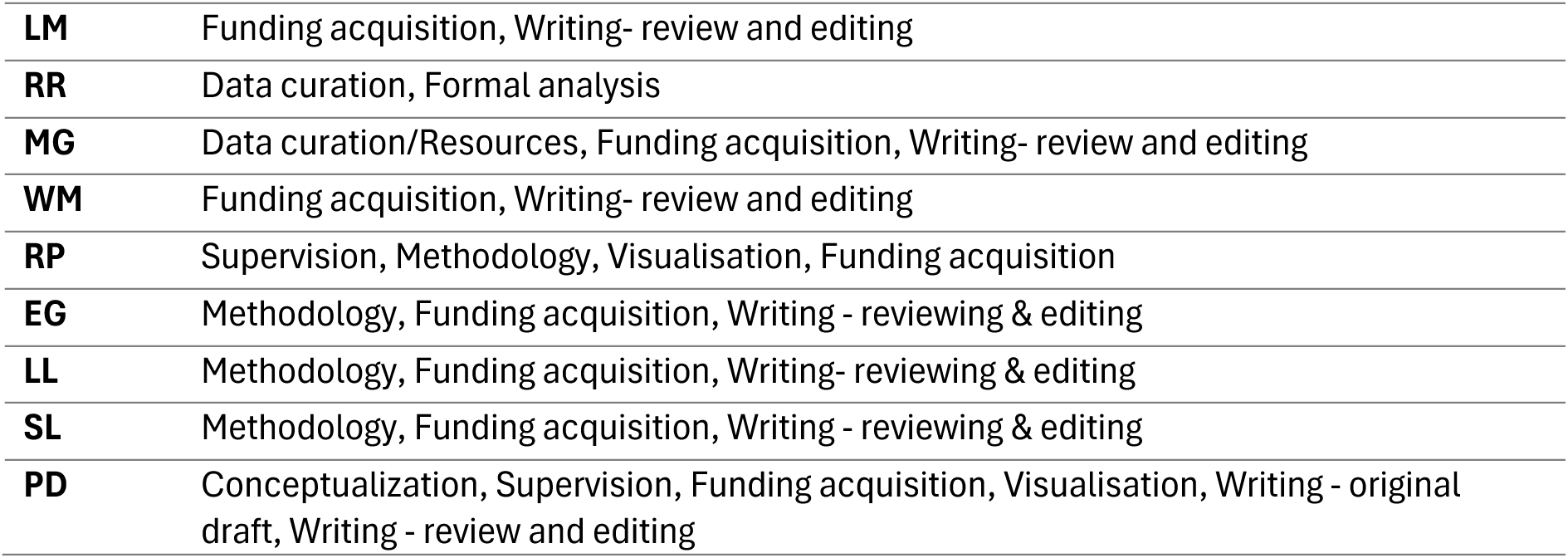

|  |  |
|---|---|
| **FDA** | Conceptualization, Data curation, Formal analysis, Investigation, Methodology, Software, Visualization, Writing - original draft, Writing - review and editing |
| **HWC** | Methodology, Software |
| **TA** | Data curation, Formal analysis, Writing- review and editing |
| **AS** | Data curation, Formal analysis |
| **LM** | Funding acquisition, Writing- review and editing |
| **RR** | Data curation, Formal analysis |
| **MG** | Data curation/Resources, Funding acquisition, Writing- review and editing |
| **WM** | Funding acquisition, Writing- review and editing |
| **RP** | Supervision, Methodology, Visualisation, Funding acquisition |
| **EG** | Methodology, Funding acquisition, Writing - reviewing & editing |
| **LL** | Methodology, Funding acquisition, Writing- reviewing & editing |
| **SL** | Methodology, Funding acquisition, Writing - reviewing & editing |
| **PD** | Conceptualization, Supervision, Funding acquisition, Visualisation, Writing - original draft, Writing - review and editing |

## Acknowledgements

This research was conducted as part of the ‘Hosts, Pathogens and Global Health’ Doctor of Philosophy programme at the University of Edinburgh (2025) and supported by a Wellcome Trust scholarship (224918/Z/21/Z) to FDA. HWC. was funded by a postgraduate studentship awarded by the Roslin Institute. RMP. is funded by a Chancellor’s Fellowship awarded by the University of Edinburgh. PD, EG, RMP, LL and SL also acknowledge Institute Strategic Programme Grant support (BBS/E/RL/230002C and BBS/E/RL/230002D) from the UK Biotechnology and Biological Sciences Research Council. WM, MG and RR acknowledge support from the South African Medical Research Council with funds received from the South African National Department of Health and the UKRI Medical Research Council, with funds received from the UK Government’s International Science Partnerships Fund (Grant No: HDID26120/KR/2024), and from the South African Research Chair initiative of the Department of Science, Technology, and Innovation, and the National Research Foundation of South Africa (Grant No. 0219114353). LM, AS and TA acknowledge funding by the UK Department for Environment, Food and Rural Affairs (Defra) and the devolved Scottish and Welsh administrations: grant numbers SE0569, SE0575, SV3700.

