## Supplementary information (Figures and smaller tables) for "The biochemical composition of coronavirus Nucleocapsid proteins drives the bicistronic properties of their sub-genomic mRNAs"

### SUPPLEMENTARY MATERIALS

**Contents:**

Supplementary Tables S1 – 4 (see separate Excel file for Table S4)

Supplementary Figures S1 – 8

Supplementary Code – Orfan script

### SUPPLEMENTARY TABLES

**Supplementary Table 1. Viral sequences used to validate ORFan.**

| Accession number | Virus isolate | Gene(s) |
|---|---|---|
| NC_002021.1 | A/Puerto Rico/8/1934(H1N1) | PB1/PB1-F2 |
| NC_004911.1 | A/Hong Kong/1073/99(H9N2) | PB1/PB1-F2 |
| NC_007358.1 | A/goose/Guangdong/1/1996 (H5N1) | PB1/PB1-F2 |
| NC_007372.1 | A/New York/392/2004(H3N2) | PB1/PB1-F2 |
| NC_007375.1 | A/Korea/426/1968(H2N2) | PB1/PB1-F2 |
| NC_026423.1 | A/Shanghai/02/2013(H7N9) | PB1/PB1-F2 |
| PX129719.1 | A/chicken/South Korea/N25-07/2025(H9N2) | PB1/PB1-F2 |
| PZ836939.1 | A/Lesser Scaup/Illinois/4297/2026(H3N6) | PB1/PB1-F2 |
| PZ837038.1 | A/Lesser Scaup/Illinois/4300/2026(H3N8) | PB1/PB1-F2 |
| PZ851521.1 | A/duck/Bangladesh/73638/2026(H10N7) | PB1/PB1-F2 |
| PZ939120.1 | A/mallard/Germany-SN/VL202603458/2026(H4N6) | PB1/PB1-F2 |
| NC_026435.1 | A/California/07/2009(H1N1) | PB1/non-functional PB1-F2 |
| GQ377049.1 | A/California/04/2009(H1N1) | PB1/non-functional PB1-F2 |
| KT873807.1 | fox-wt/ITA/288356/2011/G18P[17] | Nsp5/nsp6 |
| PZ390157.1 | RVA/Rabbit-tc/CHN/DHD01/2024/G3P[3] | Nsp5/nsp6 |
| PV631385.1 | RVA/Piglet/CHN/SQ-23/2023 | Nsp5/nsp6 |
| HQ665467.1 | Simian rotavirus A strain RRV | Nsp5/nsp6 |
| HM363564.1 | Bov/Ind/UKD/IVRI/09/M-1 | Nsp5/nsp6 |
| PV863445.1 | RVA/Horse-wt/KOR/JJ01/2024/G14P[12] | Nsp5/nsp6 |
| ON322741.1 | RVA/Dog-wt/BRA/IAL-M202/BRA/2017/G3P[3] | Nsp5/nsp6 |

| | | |
|---|---|---|
| KY055437.1 | RVA/Goat-wt/UGA/BUW-14-085/2014/G6P[1] | Nsp5/nsp6 |
| EU486985.1 | Rotavirus A isolate Ch-2 | Nsp5/nsp6 |
| LC774753.1 | RVA/Chicken/JPN/Kagoshima-840-4/2020/G19[P30] | Nsp5/truncated nsp6 |
| MZ054774.1 | RVA/Common shrew-wt/GER/KS14-269/2014/G39P[55] | Nsp5/nsp6 |
| OP718142.1 | RVA/Human-wt/MWI/BTY167/2015/G2P[4] | Nsp5/nsp6 |
| NC_077900.1 | Yaba-7 virus | N/Ns |
| NC_001927.1 | Bunyamwera virus | N/Ns |
| PX692859.1 | Oropuche virus | N/Ns |
| PV208369.1 | La Crosse virus | N/Ns |
| KP279304.1 | Schmallenberg virus | N/Ns |
| NC_009896.1 | Akabane virus | N/Ns |
| MT747974.1 | Ngari virus | N/Ns |
| MN135989.1 | Jamestown Canyon virus | N/Ns |
| AF005948.1 | Andes hantavirus | N/Ns |
| NC_005218.1 | Hantaan virus | N/ no Ns |
| Z84204.1 | Pumaala virus | N/Ns |
| MG693176.1 | CMR/Bat-AsV/P02 | CP/ORFx |
| KY855437.1 | Marmot astrovirus 1 strain HT10 | CP/truncated ORFx |
| KT946731.1 | Rodent astrovirus isolate HK-CWRN1RV | CP/ORFx |
| LC047787.1 | BoAstV/JPN/Ishikawa24-6/2013 | CP/ORFx |
| KP663426.1 | Astrovirus Er/SZAL6/HUN/2011 | CP/unknown |
| KF787112.3 | Mamastrovirus 3 isolate PAstV-GX1 | CP/ORFx |
| JN420352.1 | California sea lion astrovirus 5 isolate 1148 | CP/no ORFx |
| FJ973620.1 | Mamastrovirus 9 isolate VA1 | CP/truncated ORFx |
| Y15937.2 | Sheep astrovirus | CP/ORFx |
| EU847155.1 | Mamastrovirus 18 isolate AFCD337 | CP/ORFx |
| Z25771.1 | Human astrovirus type 1 | CP/ORFx |
| MG873154.1 | TiLV NTLV/K-2/2017 | S9/truncated S9.3 |
| OM469318.1 | NTL9/SRLAAH/2020 | S9/truncated S9.3 |
| MN939380.1 | TiLV isolate BD-2017 | S9/S9.3 |
| KY615744.1 | TiLV isolate CL | S9/S9.3 |
| MN193521.1 | TiLV WVL19031-01A | S9/S9.3 |
| ON376580.1 | TiLV HB196-VN-2020 | S9/S9.3 |
| ON376590.1 | TiLV RIA2-VN-2019 | S9/S9.3 |

| | | |
|---|---|---|
| MW814863.1 | TiLV KUTHTiLV9-2021 | uAUG S9/S9.3 |
| MN687703.1 | TiLV TH-2014 | S9/S9.3 |
| OL469281 | TiLV-2018 | uAUG S9/S9.3 |

**Supplementary Table 2. Accession number and sequence details of CoVs included in the bioinformatic analyses.** * Host species lists all known host species for each virus, not only the Refseq metadata. ^ Location lists the geographic area in which the sequence (Accession ID) was originally found.

| **Accession ID** | **Viral species** | **Genus** | **Subgenus** | **Host species*** | **Common host name** | **Date** | **Location^** | **Whole genome seq available? (Yes/No)** |
|---|---|---|---|---|---|---|---|---|
| NC_045512.2 | SARS-CoV-2 | Beta | Sarbeco | *Homo sapiens / Odocoileus virginianus* | Human, white tailed deer | 2019 | Wuhan, China | Yes |
| NC_004718.3 | SARS-CoV | Beta | Sarbeco | *Homo sapiens* | Human | 2003 | Toronto, Canada | Yes |
| AY572035 | SARS Civet CoV | Beta | Sarbeco | *Paradoxurus hermaphroditus* | Palm civet | 2004 | Guangzhou, China | Yes |
| MN996532.2 | SARS-like CoV RaTG13 | Beta | Sarbeco | *Rhinolophus affinis* | Intermediate horseshoe bat | 2013 | Yunnan, China | Yes |
| DQ084199 | SARS-like CoV HKU3-2 | Beta | Sarbeco | *Rhinolophus sinicus* | Chinese rufous horseshoe bat | 2005 | China | Yes |
| MG772933 | SARS-like CoV SL-CoVZC45 | Beta | Sarbeco | *Rhinolophus pusillus* | Least horseshoe bat | 2017 | Zhoushan, China | Yes |
| KY417146 | SARS-like CoV RS4231 | Beta | Sarbeco | *Rhinolophus sinicus* | Chinese rufous horseshoe bat | 2013 | Kunming, China | Yes |
| KF367457.1 | SARS-like CoV WIV1 | Beta | Sarbeco | *Rhinolophus sinicus* | Chinese rufous horseshoe bat | 2012 | Kunming, China | Yes |
| MT121216.1 | Pangolin/Guangdong/1/2019 CoV | Beta | Sarbeco | *Manis javanica* | Malayan pangolin | 2019 | Guandong, China | Yes |
| MT040335.1 | Pangolin/Guangxi/P5L/2007 CoV | Beta | Sarbeco | *Manis javanica* | Malayan pangolin | 2007 | Guangxi, China | Yes |

| | | | | | | | | |
|---|---|---|---|---|---|---|---|---|
| NC_014470.1 | SARS-like CoV BM48-31/BGR/2008 | Beta | Sarbeco | *Rhinolophus blasii* | Blasius's horseshoe bat | 2008 | Bulgaria | Yes |
| KJ473815.1 | SARS-like CoV GX2013 | Beta | Sarbeco | *Rhinolophus sinicus* | Chinese rufous horseshoe bat | 2013 | Guangxi, China | Yes |
| MW719567.1 | SARS-like CoV RhGB01 | Beta | Sarbeco | *Rhinolophus hipposideros* | Lesser horseshoe bat | 2020 | UK | Yes |
| KY352407.1 | SARS-like CoV BtKY72 | Beta | Sarbeco | *Rhinolophus spp. (bat)* | Horseshoe bat | 2007 | Kenya | Yes |
| OQ401252.1 | SARS-like CoV RhGB08 | Beta | Sarbeco | *Rhinolophus hipposideros* | Lesser horseshoe bat | 2020 | Falmouth, UK | Yes |
| OQ401249.1 | SARS-like CoV RfGB02 | Beta | Sarbeco | *Rhinolophus ferrumequinum* | Greater horseshoe bat | 2021 | Buckfastleigh, UK | Yes |
| OQ401248.1 | SARS-like CoV RhGB07 | Beta | Sarbeco | *Rhinolophus hipposideros* | Lesser horseshoe bat | 2020 | Falmouth, UK | Yes |
| OQ401247.1 | SARS-like CoV RfGB01 | Beta | Sarbeco | *Rhinolophus ferromequinum* | Greater horseshoe bat | 2021 | Chudleigh, UK | Yes |
| MW251308.1 | SARS-like CoV RacCS203 | Beta | Sarbeco | *Rhinolophus acuminatus* | Acuminate horseshoe bat | 2020 | Thailand | Yes |
| KT444582 | SARS-like CoV WIV16 | Beta | Sarbeco | *Rhinolophus sinicus* | Chinese rufous horseshoe bat | 2013 | Kunming, China | Yes |
| KC881005 | SARS-like CoVRsSHC014 | Beta | Sarbeco | *Rhinolophus sinicus* | Chinese rufous horseshoe bat | 2011 | China | Yes |
| KP886808 | SARS-like CoV YNLF_31C | Beta | Sarbeco | *Rhinolophus ferrumequinum* | Greater horseshoe bat | 2013 | China | Yes |
| KY938558.1 | SARS-like CoV 16BO133 | Beta | Sarbeco | *Rhinolophus ferrumequinum* | Greater horseshoe bat | 2016 | South Korea | Yes |
| MZ937000.1 | SARS-like CoV BANAL-52 | Beta | Sarbeco | *Rhinolophus malayanus* | Malayan horseshoe bat | 2020 | Feuang, Laos | Yes |

| | | | | | | | | |
|---|---|---|---|---|---|---|---|---|
| LC556375.1 | SARS-like CoV Rc-o319 | Beta | Sarbeco | *Rhinolophus cornutus* | Little Japanese horseshoe bat | 2013 | Iwate, Japan | Yes |
| MZ081381.1 | bat/Yunnan/RpYN 06/2020 | Beta | Sarbeco | *Rhinolophus pusillus* | Least horseshoe bat | 2020 | Yunnan, China | Yes |
| KF569996.1 | SARS-like CoV LYRa11 | Beta | Sarbeco | *Rhinolophus affinis* | Intermediate horseshoe bat | 2011 | Yunnan, China | Yes |
| In preparation / PX392418 | BtCoV/R.clivosus/ UP445/RWA/2008 | Beta | Sarbeco | *Rhinolophus clivosus* | Geoffroy's horseshoe bat | 2008 | Rwanda | No |
| In preparation PX392419 | BtCoV/R.blasii/UP 22-60/LP/RSA/2022 | Beta | Sarbeco | *Rhinolophus blasii* | Blasius's horseshoe bat | 2022 | South Africa | No |
| In preparation PX392420 | BtCoV/R.blasii/UP 22-65/LP/RSA/2022 | Beta | Sarbeco | *Rhinolophus blasii* | Blasius's horseshoe bat | 2022 | South Africa | No |
| In preparation PX392421 | BtCoV/R.blasii/UP 22-155/LP/RSA/2022 | Beta | Sarbeco | *Rhinolophus blasii* | Blasius's horseshoe bat | 2022 | South Africa | No |
| Unpublished PX392422 | BtCoV/R.simulator /UP22-307/LP/RSA/2022 | Beta | Sarbeco | *Rhinolophus simulator* | Bushveld horseshoe bat | 2022 | South Africa | No |
| Unpublished PX392423 | BtCoV/R.simulator /UP22-2411/LP/RSA/2022 | Beta | Sarbeco | *Rhinolophus simulator* | Bushveld horseshoe bat | 2022 | South Africa | No |
| Unpublished PX392425 | BtCoV/R.simulator /UP23-230/LP/RSA/2023 | Beta | Sarbeco | *Rhinolophus simulator* | Bushveld horseshoe bat | 2023 | South Africa | No |
| Unpublished PX392426 | BtCoV/Rhinolophus sp/BCL3/LP/RSA/2024 | Beta | Sarbeco | *Rhinolophus species* | Horseshoe bat | 2024 | South Africa | No |

| | | | | | | | | |
|---|---|---|---|---|---|---|---|---|
| Unpublished PX392427 | BtCoV/Rhinolophus sp/BCL4/LP/RSA/2024 | Beta | Sarbeco | *Rhinolophus species* | Horseshoe bat | 2024 | South Africa | No |
| Unpublished PX392424 | BtCoV/R.simulator/UP23-04/LP/RSA/2023 | Beta | Sarbeco | *Rhinolophus simulator* | Bushveld horseshoe bat | 2023 | South Africa | No |
| KJ473821.1 | BtVs-BetaCoV/SC2013 | Beta | Merbeco | *Vespertilio superans* | Asian parti-colored bat | 2013 | China | Yes |
| OQ401251.1 | Merbeco PaGB01 | Beta | Merbeco | *Plecotus auritus* | Brown long-eared bat | 2020 | Twyford, UK | Yes |
| ON378807 | MERSr-CoV B20-180 | Beta | Merbeco | *Rhinolophus ferrumequinum* | Greater horseshoe bat | 2020 | South Korea | Yes |
| NC_019843.3 | MERS-CoV | Beta | Merbeco | *Homo sapiens/ camelus dromedarius* | Human, dromedary camel | 2012 | Jeddah, Saudi Arabia | Yes |
| NC_034440.1 | MERS-related bat CoV PREDICT/PDF-2180 | Beta | Merbeco | *Pipistrellus hesperidus* | Dusky pipistrelle | 2013 | Nkuringo, Uganda | Yes |
| KC869678.4 | Neoromicia CoV PML-PHE1/RSA/2011 | Beta | Merbeco | *Neoromicia zuluensis* | Zulu serotine | 2011 | South Africa | Yes |
| KX442564.1 | Hypsugo bat CoV HKU25 | Beta | Merbeco | *Hypsugo pulveratus* | Chinese pipistrelle | 2013 | China | Yes |
| MK907286.1 | Erinaceus hedgehog corona HKU31 | Beta | Merbeco | *Erinaceus amurensis* | Amur hedgehog | 2014 | China | Yes |
| NC_039207.1 | Betacorona Erinaceus/VMC/DEU/2012 | Beta | Merbeco | *Erinaceus europaeus* | European hedgehog | 2012 | Germany | Yes |

| | | | | | | | | |
|---|---|---|---|---|---|---|---|---|
| NC_009020.1 | Pipistrellus bat CoV HKU5 | Beta | Merbeco | *Pipistrellus abramus* | Japanese house bat | 2005 | Guandong, China | Yes |
| NC_009019.1 | Tylonycteris bat CoV HKU4 | Beta | Merbeco | *Tylonycteris pachypus* | Lesser bamboo bat | 2006 | Guandong, China | Yes |
| MN514964.1 | BetaCoV HKU23 strain HKU23/camel/Nigeria/NV1010/2015 | Beta | Embeco | *Camelus dromedarius* | Dromedary camel | 2015 | Nigeria | Yes |
| NC_006577.2 | Human CoV HKU1 | Beta | Embeco | *Homo sapiens* | Human | 2004 | Shenzen, China | Yes |
| NC_001846.1 | MHV | Beta | Embeco | *Mus musculus* | Common mouse | 1949 | USA | Yes |
| EF446615.1 | Equine CoV NC99 | Beta | Embeco | *Equus ferus caballus* | Horse | 2000 | USA | Yes |
| NC_003045.1 | Bovine CoV | Beta | Embeco | *Bos taurus* | Cattle | 2000 | USA | Yes |
| NC_046954.1 | Rodent CoV RtMruf-CoV-2/JL2014 | Beta | Embeco | *Myodes rufocanus* | Grey red-backed vole | 2014 | China | Yes |
| NC_012936.1 | Rat Corona Parker | Beta | Embeco | *Rattus Rattus* | Black rat | 2009 | USA | Yes |
| NC_026011.1 | China Rattus CoV HKU24 | Beta | Embeco | *Rattus norvegicus* | Brown rat | 2012 | China | Yes |
| NC_006213.1 | Human CoV OC43 | Beta | Embeco | *Homo sapiens* | Human | 2004 | USA | Yes |
| DQ011855 | PHEV | Beta | Embeco | *Sus scrofa domesticus* | Pig | 1972 | Belgium | Yes |
| EF424623.1 | Giraffe CoV strain US/OH3/2003 | Beta | Embeco | *Giraffa camelopardalis* | Giraffe | 2003 | USA | Yes |
| NC_017083.1 | Rabbit CoV HKU14 | Beta | Embeco | *Oryctolagus cuniculus* | European rabbit | 2006 | China | Yes |
| MZ100070.1 | Tapir CoV | Beta | Embeco | *Acrocodia indica* | Indonesian tapir | 2021 | Canada | Yes |

| | | | | | | | | |
|---|---|---|---|---|---|---|---|---|
| OM397541.1 | Oryx CoV | Beta | Embeco | *Oryx leucoryx* | Arabian Oryx | 2019 | Israel | Yes |
| NC_009021.1 | Rousettus bat HKU9 | Beta | Nobeco | *Rousettus spp.* | Rousette bat | 2006 | Guandong, China | Yes |
| NC_030886.1 | Rousettus bat CoV GCCDC1 | Beta | Nobeco | *Rousettus leschenaulti* | Leschenault's rousette | 2014 | Yunnan, China | Yes |
| NC_048212.1 | Eidolon Helvum Bat betacorona | Beta | Nobeco | *Eidolon Helvum* | Straw-coloured fruit bat | 2013 | Cameroon | Yes |
| HQ728482.1 | Eidolon bat corona/Kenya/KY 24/2006 | Beta | Nobeco | *Eidolon Helvum* | Straw-coloured fruit bat | 2006 | Kenya | Yes |
| OK067319.1 | P.rufus CoV AMB130 | Beta | Nobeco | *Pteropus rufus* | Madagascan flying fox | 2018 | Madagascar | Yes |
| PQ179293.1 | NobeCoV NRB24 | Beta | Nobeco | *Epomophorus wahlbergi* | Wahlberg's epauletted fruit bat | 2023 | Kenya | Yes |
| PX392226 | BtCoV_UPE1000 | Beta | Nobeco | *Rousettus aegyptiacus* | Egyptian fruit bat | 2024 | South Africa | No |
| HQ166910.1 | Zaria bat CoV | Beta | Hibeco | *Hipposideros commersoni* | Commerson's leaf-nosed bats | 2008 | Nigeria | Yes |
| NC_025217.1 | Bat Hp CoV Zhejiang2013 | Beta | Hibeco | *Hipposideros pratti* | Pratt's roundleaf bat | 2013 | Zhejiang, China | Yes |
| OQ469322 | Hipposideros armiger Corona isolate CD36 | Beta | Hibeco | *Hipposideros armiger* | Great roundleaf bat | 2021 | Hainan, China | Yes |
| OK017908 | Betacorona sp. isolate CD35 | Beta | Hibeco | *Hipposideros armiger* | Great roundleaf bat | 2021 | Hainan, China | Yes |
| NC_010800.1 | Turkey CoV | Gamma | Igaco | *Meleagris gallopavo* | Wild turkey | 2007 | Canada | Yes |
| NC_048213.1 | IBV | Gamma | Igaco | *Gallus gallus* | Chicken | 2003 | India | Yes |

| | | | | | | | | |
|---|---|---|---|---|---|---|---|---|
| NC_048214.1 | Duck CoV | Gamma | Igaco | *Aix galericulata* | Mandarin duck | 2014 | Guandong, China | Yes |
| AJ871017.1 | Goose CoV_03/586-50 | Gamma | Brangaco | *Anser anser* | Graylag goose | 2003 | Norway | No |
| MW436465.1 | Anser fabalis CoV NCN2 | Gamma | Brangaco | *Anser fabalis* | Taiga veab goose | 2019 | China | Yes |
| MW588092.1 | Mute swan CoV Atonsbury/A/2016 | Gamma | Brangaco | *Cygnus olor* | Mute swan | 2016 | UK | Yes |
| NC_046965.1 | Canada Goose CoV | Gamma | Brangaco | *Branta canadensis* | Canada goose | 2017 | Canada | Yes |
| NC_010646.1 | Beluga whale CoV SW1 | Gamma | Cegaco | *Delphinapterus leucas* | Beluga whale | 2007 | USA | Yes |
| KF793826.1 | Bottlenose dolphin CoV HKU22 | Gamma | Cegaco | *Tursiops aduncus* | Bottlenose dolphin | 2010 | Hong Kong | Yes |
| NC_016995.1 | Wigeon CoV HKU20 | Delta | Andeco | *Mareca penelope* | Eurasian wigeon | 2008 | Hong Kong | Yes |
| NC_016994.1 | Night heron CoV HKU19 | Delta | Herdeco | *Nycticorax nycticorax* | black-crowned night heron | 2007 | Hong Kong | Yes |
| NC_016996.1 | Common moorhen CoV HKU21 | Delta | Buldeco | *Gallinula chloropus* | Common moorhen | 2007 | Hong Kong | Yes |
| LC364342.1 | Falcon CoV HKU27 | Delta | Buldeco | *Falco spp.* | Falcon | 2013 | UAE | Yes |
| NC_011550.1 | Munia CoV HKU13-3514 | Delta | Buldeco | *Lonchura striata* | White-rumped munia | 2007 | Hong Kong | Yes |
| NC_016993.1 | Magpie-robin CoV HKU18 | Delta | Buldeco | *Copsychus saularis* | Oriental magpie-robin | 2007 | Hong Kong | Yes |
| LC364343.1 | Houbara CoV HKU28 | Delta | Buldeco | *Chlamydotis sp.* | Houbara bustard species | 2013 | UAE | Yes |

| | | | | | | | | |
|---|---|---|---|---|---|---|---|---|
| NC_016992.1 | Sparrow CoV HKU17 | Delta | Buldeco | *Passer montanus* | Eurasian tree sparrow | 2007 | Hong Kong | Yes |
| LC364344.1 | Pigeon CoV HKU29 | Delta | Buldeco | *Columba spp.* | Pidgeon species | 2013 | UAE | Yes |
| NC_039208.1 | PDCoV | Delta | Buldeco | *Sus scrofa domesticus* | Pig | 2010 | Hong Kong | Yes |
| NC_016991.1 | White-eye CoV HKU16 | Delta | Buldeco | *Zosterops spp.* | Passerine bird | 2007 | Hong Kong | Yes |
| NC_011549.1 | Thrush CoV HKU12-600 | Delta | Buldeco | *Turdus hortulorum* | Grey-backed thrush | 2007 | Hong Kong | Yes |
| NC_011547.1 | Bulbul CoV HKU11-934 | Delta | Buldeco | *Pycnonotus jocosus* | Red-whiskered bulbul | 2007 | Hong Kong | Yes |
| LC364346.1 | Quail CoV UAE-HKU30 | Delta | Buldeco | *Coturnix coturnix* | Common quail | 2013 | UAE | Yes |
| PP397109.1 | Black-headed gull CoV | Delta | Buldeco | *Chroicocephalus ridibundus* | Black-headed gull | 2021 | China | Yes |
| NC_046955.1 | Shrew CoV | Alpha | Sunaco | *Sorex araneus* | Common shrew | 2014 | Tibet | Yes |
| MZ328303.1 | Wenzhou Suncus murinus corona | Alpha | Sunaco | *Suncus murinus* | Asian house shrew | 2016 | Wenzhou, China | Yes |
| NC_048211.1 | Wencheng Sm Shrew CoV | Alpha | Sunaco | *Suncus murinus* | Asian house shrew | 2015 | Wencheng, China | Yes |
| NC_034972 | Luchaco AcCoV-JC34 | Alpha | Luchaco | *Apodemus chevrieri* | Chevrier's field mouse | 2011 | Yunna, China | Yes |
| NC_032730.1 | Lucheng Rn Rat CoV | Alpha | Luchaco | *Rattus norvegicus* | Grey rat | 2013 | Zhejiang, China | Yes |
| PP868300.1 | HCoV KUMC22-3 | Alpha | Luchaco | *Homo sapiens* | Human | 2022 | South Korea | Yes |
| MZ328301.1 | Berylmys bowersi alphaCoV-1 | Alpha | Luchaco | *Berylmys bowersi* | Bower's white-toothed rat | 2016 | China | Yes |

| | | | | | | | | |
|---|---|---|---|---|---|---|---|---|
| MZ577265.1 | Plateatu pika CoV P83 | Alpha | Luchaco | *Ochotona curzoniae* | Plateau pika | 2021 | China | Yes |
| NC_038861.1 | TGEV | Alpha | Tegaco | *Sus scrofa domestica* | Pig | 2000 | USA | Yes |
| PP069800.1 | SADS-CoV | Alpha | Rhinaco | *Sus scrofa domestica* | Pig | 2023 | China | Yes |
| MW591993.2 | Canine CoV CCoV-HuPn2018 | Alpha | Tegaco | *Homo sapiens* | Human | 2018 | Malaysia | Yes |
| NC_002306.3 | FIPV | Alpha | Tegaco | *Felis catus* | Common cat | 2010 | USA | Yes |
| NC_023760.1 | Mink CoV WD1127 | Alpha | Minaco | *Mustela vison* | American mink | 1998 | USA | Yes |
| NC_030292.1 | Ferret alphaCoV | Alpha | Minaco | *Mustela putorius* | European polecat | 2010 | Netherlands | Yes |
| NC_002645.1 | Human CoV 229E | Alpha | Duvinaco | *Homo sapiens* | Human | 2000 | Germany | Yes |
| NC_028752.1 | Camel alphaCoV | Alpha | Duvinaco | *Camelus bactrianus* | Mongolian camel | 2015 | Riyadh, Saudi Arabia | Yes |
| KY073747.1 | 229E-related CoV BtKY229E1 | Alpha | Duvinaco | *Hipposideros spp.* | Round leaf bat | 2009 | Kenya | Yes |
| KY073748.1 | 229E-related CoV BtKY229E8 | Alpha | Duvinaco | *Hipposideros vittatus* | Striped leaf-nosed bat | 2010 | Kenya | Yes |
| NC_028824.1 | RhinaCoV BtRfAlphaCoV/YN 2012 | Alpha | Rhinaco | *Rhinolophus ferrumequinum* | Greater horseshoe bat | 2012 | China | Yes |
| NC_009988.1 | Rhinolophus bat CoV HKU2 | Alpha | Rhinaco | *Rhinolophus sinicus* | Chinese rufous horseshoe bat | 2006 | Guandong, China | Yes |
| OQ175250.1 | Bat Coronavirus RsHI19 | Alpha | Rhinaco | *Rhinolophus sinicus* | Chinese rufous horseshoe bat | 2019 | Hainan, China | Yes |
| PP746500.1 | Rhinacovirus sp. Rt22CB395R | Alpha | Rhinaco | *Rhinolophus thomasi* | Thomas's horseshoe bat | 2022 | Vietnam | Yes |

| | | | | | | | | |
|---|---|---|---|---|---|---|---|---|
| NC_048216.1 | NL63-related bat CoV BtKYNL63-9b | Alpha | Setraco | *Triaenops afer* | African trident bat | 2010 | Kenya | Yes |
| NC_032107.1 | NL63-related bat CoV BtKYNL63-9a | Alpha | Setraco | *Triaenops afer* | African trident bat | 2010 | Kenya | Yes |
| KY073746.1 | NL63-related bat CoV BtKYNL63-15 | Alpha | Setraco | *Triaenops afer* | African trident bat | 2008 | Kenya | Yes |
| NC_005831.2 | Human CoV NL63 | Alpha | Setraco | *Homo sapiens* | Human | 2004 | Netherlands | Yes |
| NC_009657.1 | Scotophilus bat CoV 512 | Alpha | Pedaco | *Scotophilus kuhlii* | Lesser Asiatic yellow bat | 2005 | China | Yes |
| NC_003436.1 | PEDV | Alpha | Pedaco | *Sus scrofa domestica* | Pig | 1993 | Belgium | Yes |
| OR248861.1 | Pekapeka alphaCoV | Alpha | Pedaco | *Chalinolobus tuberculatus* | New Zealand long-tailed bat | 2020 | New Zealand | Yes |
| unpublished | Alphacov/M.dau/APHA/2025 | Alpha | Pedaco | *Myotis daubentonii* | Daubenton's bat | 2025 | UK | No |
| unpublished | Alphacov/M.nat/APHA/2025 | Alpha | Pedaco | *Myotis nattereri* | Natterer's bat | 2025 | UK | No |
| OQ401255.1 | Myotis CoV MdGB03 | Alpha | Pedaco | *Myotis daubentonii* | Daubenton's bat | 2020 | UK | Yes |
| OQ401254.1 | Myotis CoV MdGB02 | Alpha | Pedaco | *Myotis daubentonii* | Daubenton's bat | 2020 | UK | Yes |
| OQ401253.1 | Pipistrellus alphaCoV PpiGB01 | Alpha | Pedaco | *Pipistrellus pipistrellus* | Common pipistrelle | 2021 | UK | Yes |
| OQ401250.1 | Myotis CoV MdGB01 | Alpha | Pedaco | *Myotis daubentonii* | Daubenton's bat | 2020 | UK | Yes |

| | | | | | | | | |
|---|---|---|---|---|---|---|---|---|
| NC_022103.1 | Bat CoV CDPHE15/USA/2006 | Alpha | Cola | *Myotis lucifugus* | Little brown bat | 2006 | USA | Yes |
| NC_028811.1 | M.ricketti CoV Sax-2011 | Alpha | Myotaco | *Myotis ricketti* | Rickett's big-footed bat | 2011 | China | Yes |
| MW924112 | Alphacoronavirus HCQD-2020 | Alpha | Myotaco | *Eptescus serotinus* | Serotine bat | 2020 | Korea | Yes |
| MZ328298.1 | M.chinensis alphacoronavirus | Alpha | Myotaco | *Myotis chinensis* | Large vesper bat | 2016 | China | Yes |
| NC_028833.1 | Nyctalus velutinus alphacorona SC-2013 | Alpha | Nyctaco | *Nyctalus velutinus* | Chinese noctule | 2013 | China | Yes |
| NC_046964.1 | Alphacorona BatCoV/P.kuhlii/Italy/339819/2015 | Alpha | Nyctaco | *Pipistrellus kuhlii* | Kuhl’s pipistrelle | 2015 | Italy | Yes |
| NC_054015.1 | T.bat CoV HKU33-GZ151867 | Alpha | Nyctaco | *Tylonycteris robustula* | Greater bamboo bat | 2015 | China | Yes |
| Unpublished | Aphacov/E.ser/Bat24-1156 | Alpha | Nyctaco | *Eptescus serotinus* | Serotine bat | 2024 | UK | No |
| NC_010438.1 | Miniopterus bat CoV HKU8 | Alpha | Minunaco | *Miniopterus spp.* | Long winged bat | 2004 | Hong Kong | Yes |
| NC_010437.1 | Miniopterus bat CoV 1A | Alpha | Minunaco | *Miniopterus magnater* | Western bent-winged bat | 2004 | Hong Kong | Yes |
| MK472070.1 | AlphaCoV WA3607 | Alpha | Decaco | *Ozimops spp.* | Molossid bat | 2018 | Australia | Yes |
| NC_028814.1 | R.ferrumequinum alphaCoV HuB-2013 | Alpha | Decaco | *Rhinolophus ferrumequinum* | Greater horseshoe bat | 2013 | China | Yes |
| NC_018871.1 | Rousettus bat CoV HKU10 | Alpha | Decaco | *Rousettus leschenaultii* | Leschenault's rousette fruit bat | 2005 | China | Yes |

| | | | | | | | | |
|---|---|---|---|---|---|---|---|---|
| MN611525.1 | H.pomona bat CoV CHB25 | Alpha | Decaco | *Hipposideros pomona* | Pomona roundleaf bat | 2018 | China | Yes |
| OK287352.1 | Yunnan/Hp_JC8xc/2020 | Alpha | Decaco | *Hipposideros pomona* | Pomona roundleaf bat | 2020 | China | Yes |
| OK287353.1 | Yunnan/Rs/CX18c/2020 | Alpha | Decaco | *Rhinolophus sinicus* | Chinese rufous horseshoe bat | 2020 | China | Yes |
| OK017907.1 | H.pomona CoV sp. isolate CD30 | Alpha | Decaco | *Hipposideros pomona* | Pomona roundleaf bat | 2021 | China | Yes |
| PQ649435.1 | BtCoV/Rf/Russia/F3/2020 | Alpha | Decaco | *Rhinolophus ferrumequinum* | Greater horseshoe bat | 2020 | Russia | Yes |
| MK720945.1 | R.bat CoV HKU32-TLC26A | Alpha | Decaco | *Rhinolophus sinicus* | Chinese rufous horseshoe bat | 2015 | Hong Kong | Yes |
| NC_055953.1 | Bat CoV AMA_L_F | Alpha | Unknown (tentative Soracovirus) | *Desmodus rotundus* | Common vampire bat | 2015 | Peru | Yes |

**Supplementary Table 3. Predicted TRS-L and B sequences, divided by subgenus and structural gene.**

| **Genus** | **Subgenus** | **Putative TRS-L** | **Putative TRS-B-N** | **Putative TRS-B-M** | **Putative TRS-B-E** | **Putative TRS-B-S** |
|---|---|---|---|---|---|---|
| Alpha | Colavirus | ATCTAAC | AACTAAAC | AACGAAA | AACTCGAC | TACTAA |
| | Decacovirus | AACTAAAC | AACTAAAC | AACTAAAC | AACTAAAC | AACTAAAC |
| | Duvinacovirus | AACTAAA | ATCTAAA | ATCTAAA | ATCTAAA | ATCTAAA |
| | Luchacovirus | AACTAAAC | AACTAAA | AACTAAA | AACTAAA | AACTAAA |
| | Minacovirus | AACTAAAC | AACTAAAC | AACTAAAC | AACTAAAC | AACTAAAC |
| | Minunacovirus | AACTAAAC | AACTAAAC | ATCTAAAC | AACTACAC | AACTAAA |

| | | | | | | |
|---|---|---|---|---|---|---|
| | Myotacovirus | AACTAAAC | AACTAAAC | AACTAAA | AACTTGA | AACTAAA |
| | Nyctacovirus | AACTGCAAC | ACCGAAAC | AACGAAA | AACTAAAC | AACTAAAT |
| | Pedacovirus | AACTAAAC | GTCTAAAC | AATCATTT | AACTAGAC | TCTAATC |
| | Rhinacovirus | AACTAAAC | AACTAAAC | AACTAAAC | AACTAAAC | AACTAAAT |
| | Setracovirus | AACTAAAC | AACTAAAC | AACTAAAC | AACTATAC | AACTAAAT/C |
| | Soracovirus | AACTAAAC | AACTAAAC | AACCAAA | AACTAAAC | AACTAA |
| | Sunacovirus | AACTAAAC | GCCTAAAC | AACTAAAC | TACTAAAC | AACTAAC |
| | Tegacovirus | AACTAAAC | AACTAAAC | AACTAAAC | TATAAAC | ACTAAAC |
| Beta | Embecovirus | ATCTAAAC | ATCTAAAC | ATCCAAAC | ATCCAAAC | ATCTAAAC |
| | Hibecovirus | ACGAAC | ACGAAC | ACGAAC | ACGAAC | ACGGAC |
| | Merbecovirus | ACGAAC | ACGAATC | ACGAAC | ACGAAC | ACGAAC |
| | Nobecovirus | ACGAAC | ACGAAC | ACGAAC | TCGAAC | ACGAAC |
| | Sarbecovirus | ACGAAC | ACGAAC | ACGAAC | ACGAAC | ACGAAC |
| Gamma | Brangacovirus | CTTAACA | CTTAACA | TTAACA | CTTAACC | CTTAAAA |
| | Cegacovirus | ATTAAAC | ATTACAAC | ATTAAAC | AATAAA | ATTTAAA |
| | Igacovirus | CTTAACA | CTTAACA | CTTAACA | CTAACA | CTGAAC |
| Delta | Andecovirus | GACACCA | AACACCA | GACACC | CAATACCA | AAAACCAA |
| | Buldecovirus | GACACCA | GACACCA | ACACCA | ACACCA | GACACCA |
| | Herdecovirus | GACACCGA | GACACCGA | GGCACCA | GACACCAA | GCCACCCA |

21

### SUPPLEMENTARY FIGURES

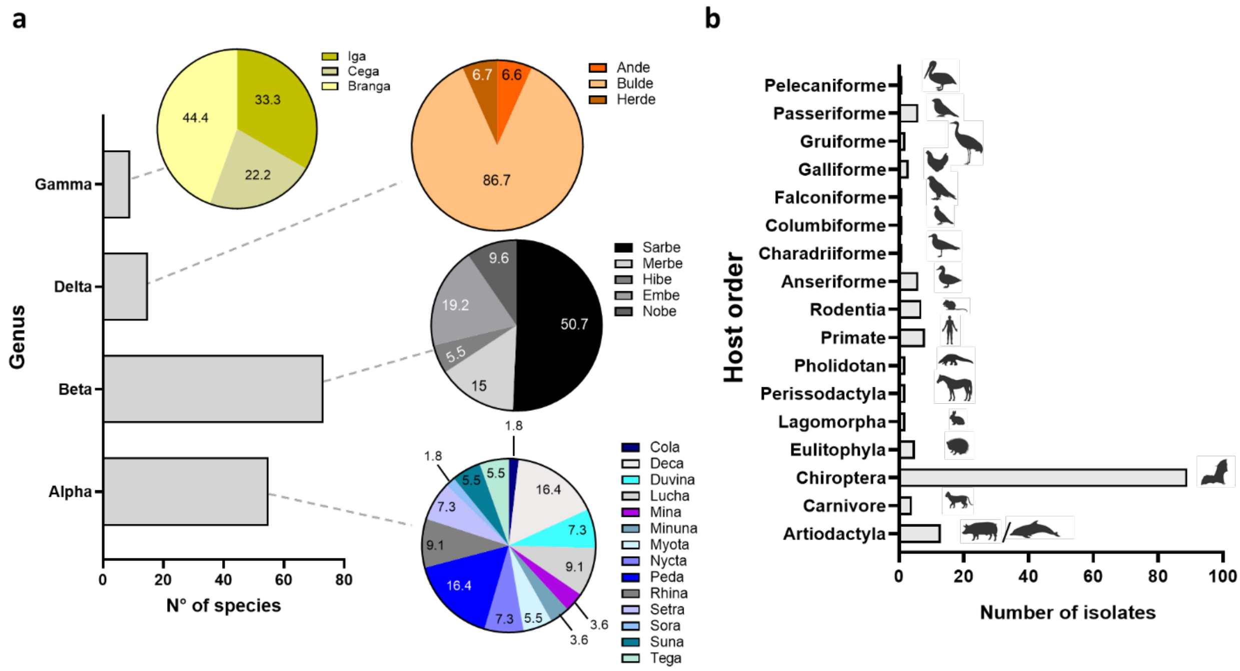


**Supplementary figure 1. CoV N sequence data distribution by genus, subgenus and host organism.** a) Number of viral species divided by genus with subgenus percentage breakdowns. b) Sequence distribution by host order.

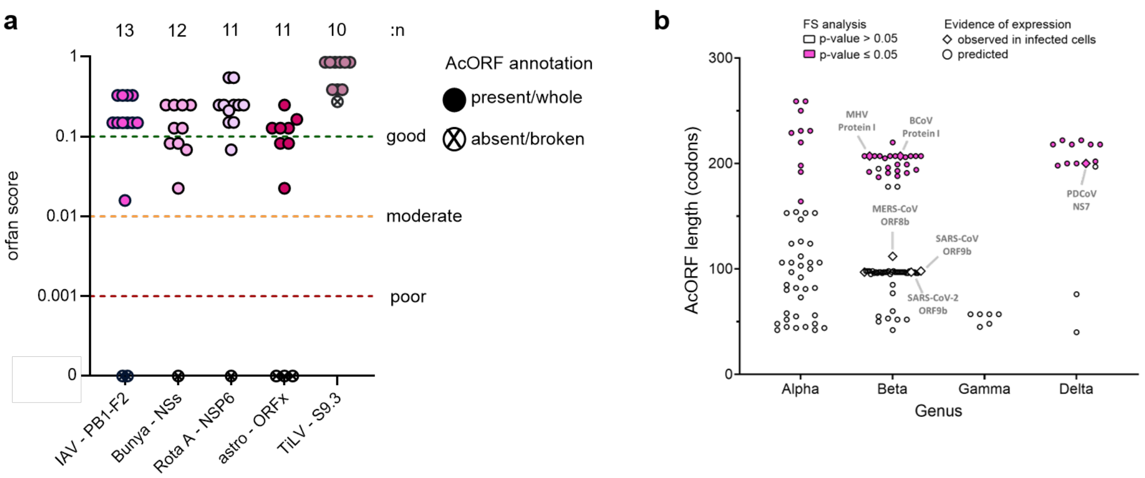


**Supplementary Figure 2. ORFan sensitivity and specificity evaluation**. a) Individual AcORF likelihood score divided by viral family and AcORF species. An ORFan score ≥0.1 suggests a good (green line) accessory gene candidate**.** 0.1>x≥0.01 a moderate (yellow line), 0.01>x≥0.001 poor (red line) and <0.001 was considered not to have a candidate. Numbers above indicate the number of sequences analysed for each virus family/species. b) Dot plot of CoV AcORF candidate lengths (in codons) identified by ORFan. AcORFs encoding an experimentally confirmed polypeptide are shown as diamonds. AcORF candidates classified by as statistically significant by the FS analysis are shown in fuchsia.

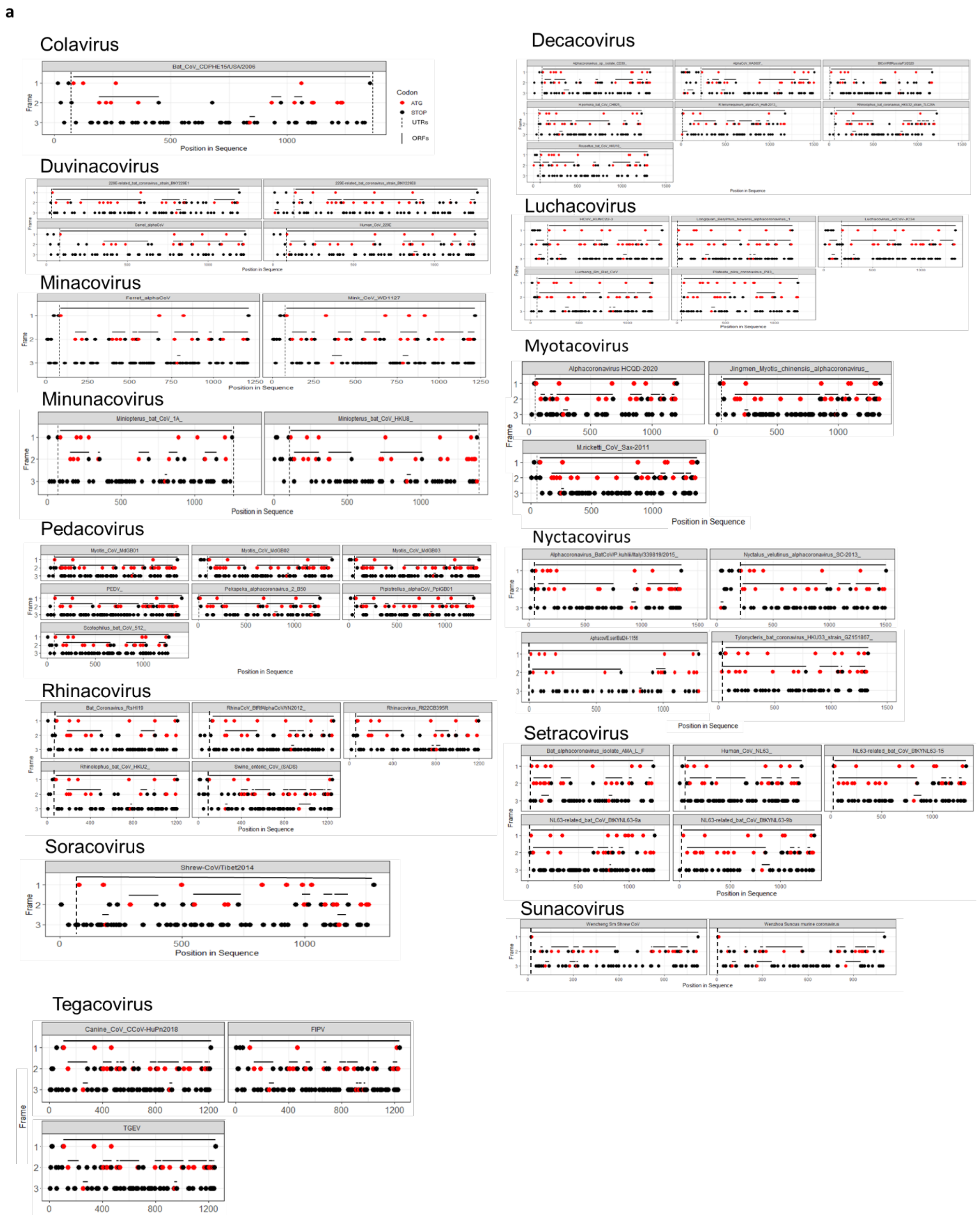
a
Colavirus
Bat_CoV_CDPHE15/USA/2006
Codon
ATG
STOP
UTRs
ORFs
Position in Sequence
Frame
Duvinacovirus
Minacovirus
Ferret_alphaCoV
Mink_CoV_WD1127
Minunacovirus
Miniopterus_bat_CoV_1A_
Miniopterus_bat_CoV_HKU8_
Pedacovirus
Rhinacovirus
Soracovirus
Shrew-CoV/Tibet2014
Tegacovirus
Canine_CoV_CCoV-HuPn2018
FIPV
TGEV
Decacovirus
Luchacovirus
Myotacovirus
Alphacoronavirus HCQD-2020
Jingmen_Myotis_chinensis_alphacoronavirus_
M.ricketti_CoV_Sax-2011
Nyctacovirus
Setracovirus
Bat_alphacoronavirus_isolate_AMA_L_F
Human_CoV_NL63_
NL63-related_bat_CoV_BtKYNL63-15
NL63-related_bat_CoV_BtKYNL63-9a
NL63-related_bat_CoV_BtKYNL63-9b
Sunacovirus

b

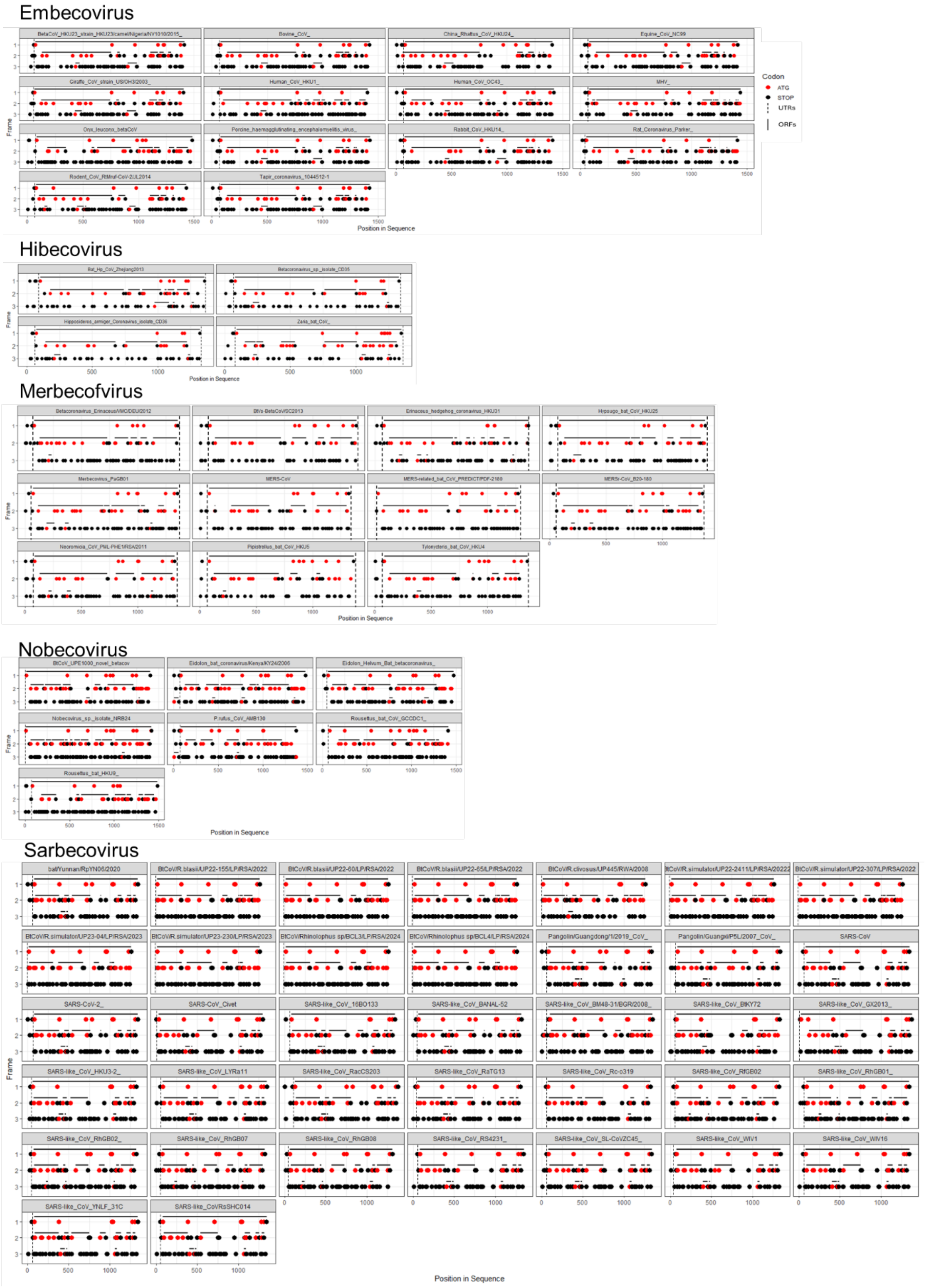
Embecovirus
Hibecovirus
Merbecofvirus
Nobecovirus
Sarbecovirus
Codon
ATG
STOP
UTRs
ORFs
Frame
Position in Sequence
bat/Yunnan/RpYN06/2020
SARS-CoV-2_
SARS-CoV_Civet
SARS-CoV
SARS-like_CoV_16BO133
SARS-like_CoV_BANAL-52
SARS-like_CoV_HKU3-2_
SARS-like_CoV_LYRa11
SARS-like_CoV_RacCS203
SARS-like_CoV_RaTG13
SARS-like_CoV_Rc-o319
SARS-like_CoV_RfGB02
SARS-like_CoV_RhGB01_
SARS-like_CoV_RhGB02_
SARS-like_CoV_RhGB07
SARS-like_CoV_RhGB08
SARS-like_CoV_RS4231_
SARS-like_CoV_SL-CoVZC45_
SARS-like_CoV_WIV1
SARS-like_CoV_WIV16
SARS-like_CoV_YNLF_31C
SARS-like_CoVRsSHC014

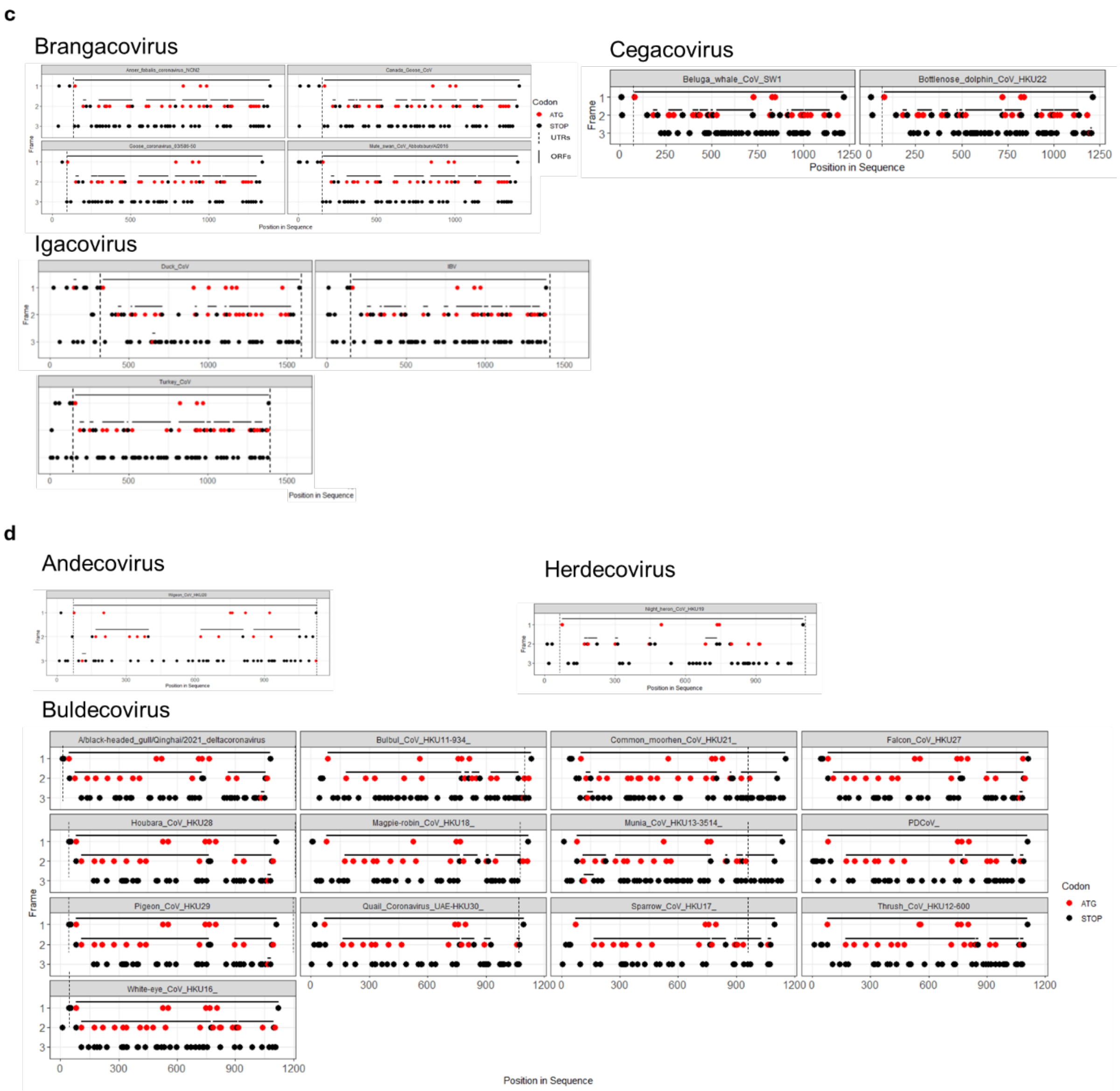


**Supplementary figure 3. ORF distribution across the three reading frames of CoV N genes. Plots show** ATG (red spots) and Stop codons (black spots) across positive sense reading frames within the N sgmRNA molecules of a) α-, b) β-, c) γ- and d) δ-CoVs. Each box represents a different sequence/Cov species. Solid lines represent ORFs. Dashed lines delimit the 5'UTRs of each sgmRNA.

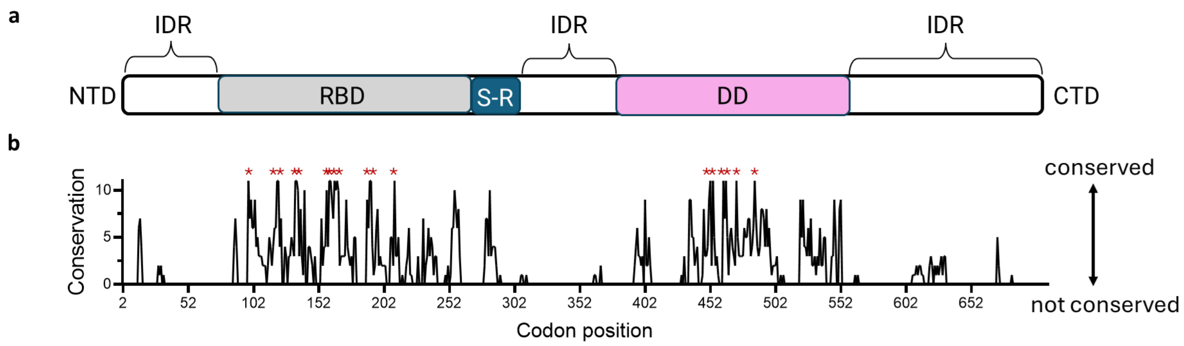


**Supplementary figure 4. Spatial conservation of N functional domains.** a) Linear diagram of the Nucleocapsid protein structure (RNA-binding domain (RBD, grey); dimerization domain (DD, fuchsia); serine-arginine (S-R) linker, blue; intrinsically disordered regions (IDRs, white). b) Residue-by-residue biochemical conservation score of the N protein across the CoV family, generated using Jalview. Residues conserved across the whole CoV dataset are highlighted with red asterisks.

**a**

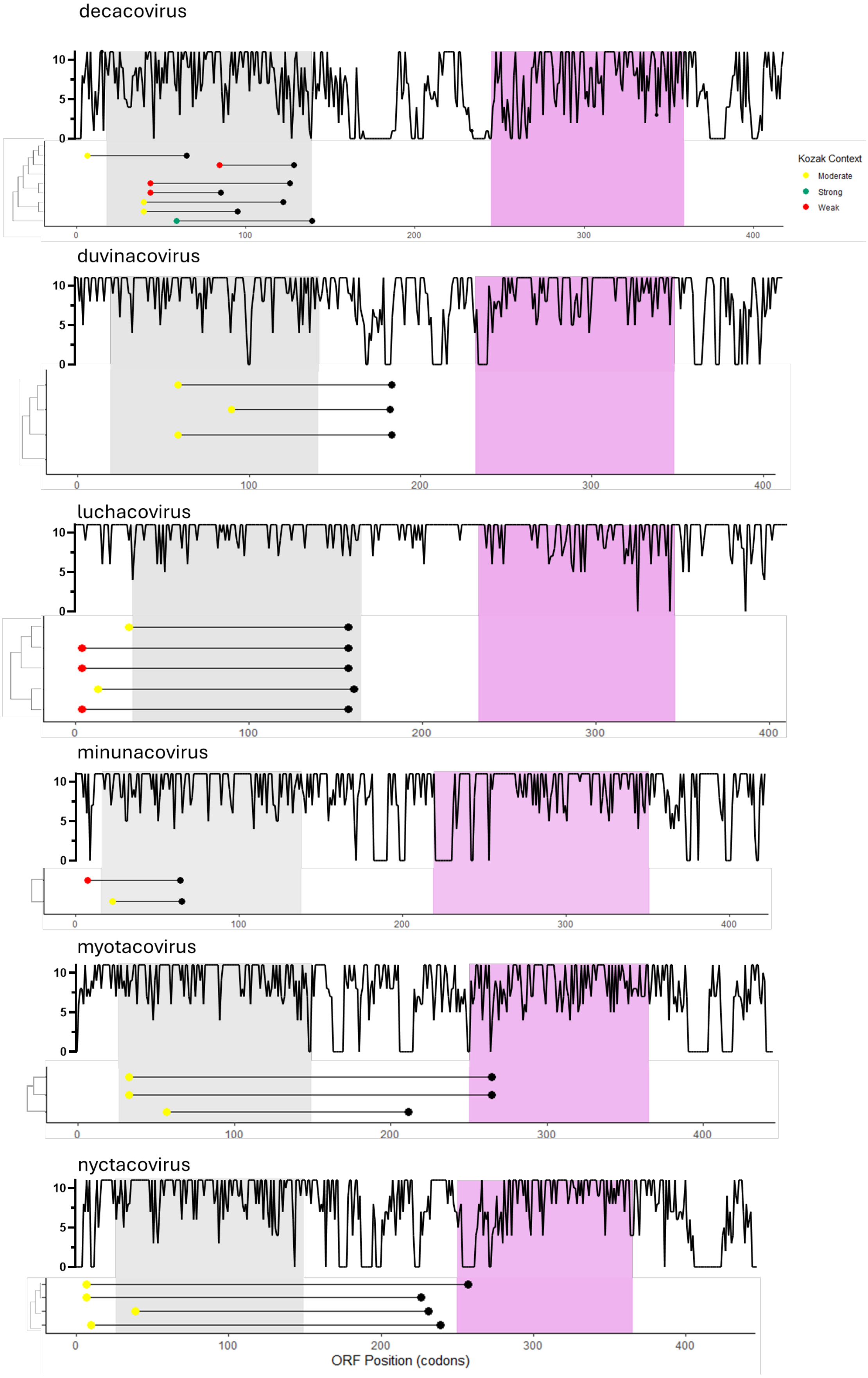
decacovirus
Kozak Context
Moderate
Strong
Weak
duvinacovirus
luchacovirus
minunacovirus
myotacovirus
nyctacovirus
ORF Position (codons)

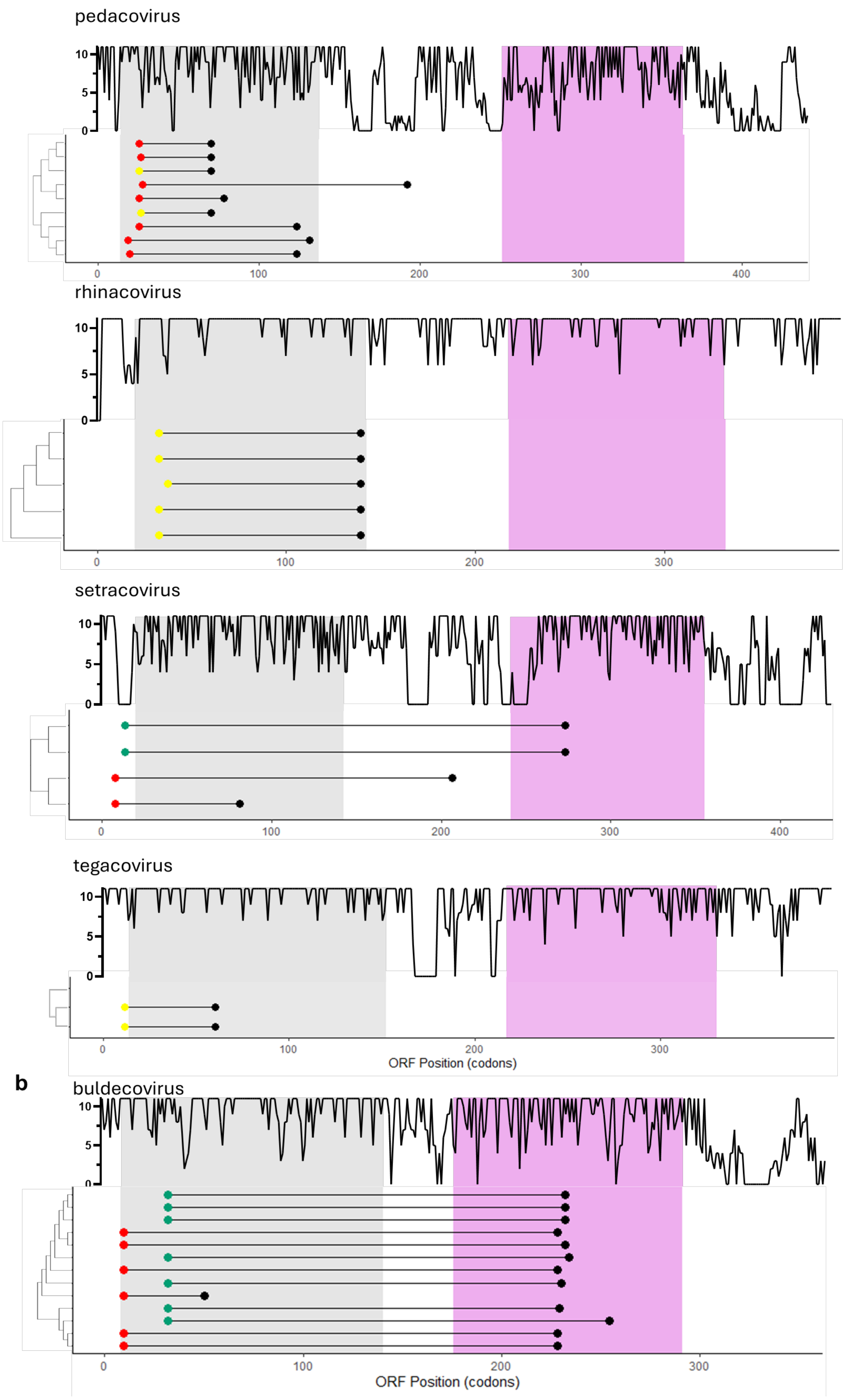
pedacovirus
10
5
0
0
100
200
300
400
rhinacovirus
10
5
0
0
100
200
300
setracovirus
10
5
0
0
100
200
300
400
tegacovirus
10
5
0
0
100
200
300
ORF Position (codons)
b
buldecovirus
10
5
0
0
100
200
300
ORF Position (codons)

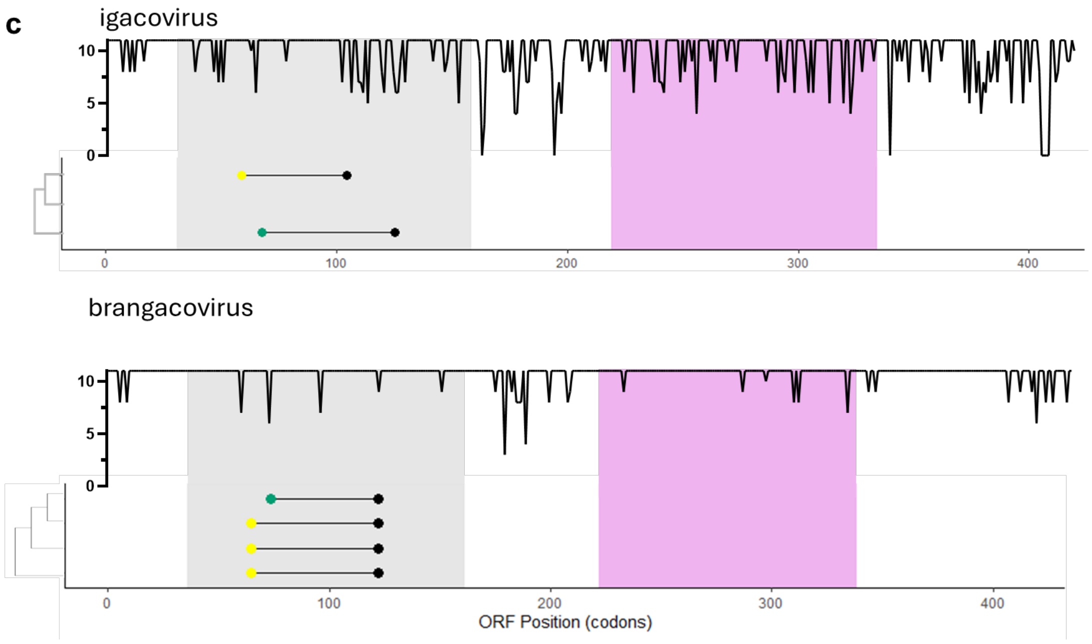


**Supplementary figure 5. N protein conservation and AcORF genomic coordinates in α-, δ- and γ-CoVs.** N protein residue-by-residue biochemical conservation profile and species-specific frame 2 AcORF codon coordinates for a) α-, b) δ- and c) γ-CoVs, divided by subgenus. The median coordinates of N RBD (grey) and DD (fuchsia) regions and the codon coordinates of each Frame 2 AcORF were plotted using RStudio. Cladograms were generated using MEGA11. Start codons of AcORFs and N genes are coloured according to their Kozak context (green = strong; yellow = moderate; red = weak).

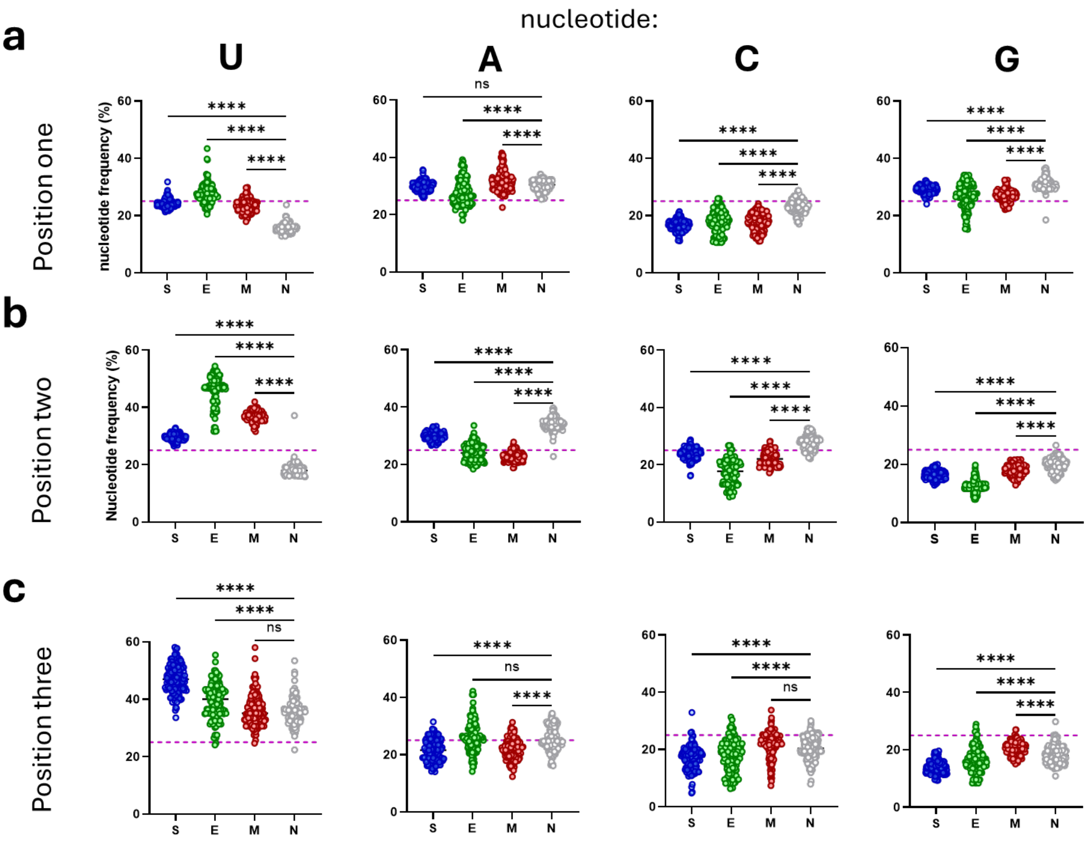


**Supplementary figure 6. Nucleotide compositional biases of CoV structural genes.** Average nucleotide distribution at a) position one; b) two and c) three of each codon in CoV structural genes. Each dot represents the average nucleotide distribution within an individual sequence. Statistical significance was calculated via One-Way ANOVA with the Bonferroni correction for multiple comparisons (ns: non-significant, *: p<0.05; **: p<0.01; ***: p<0.001; ****: p<0.0001).

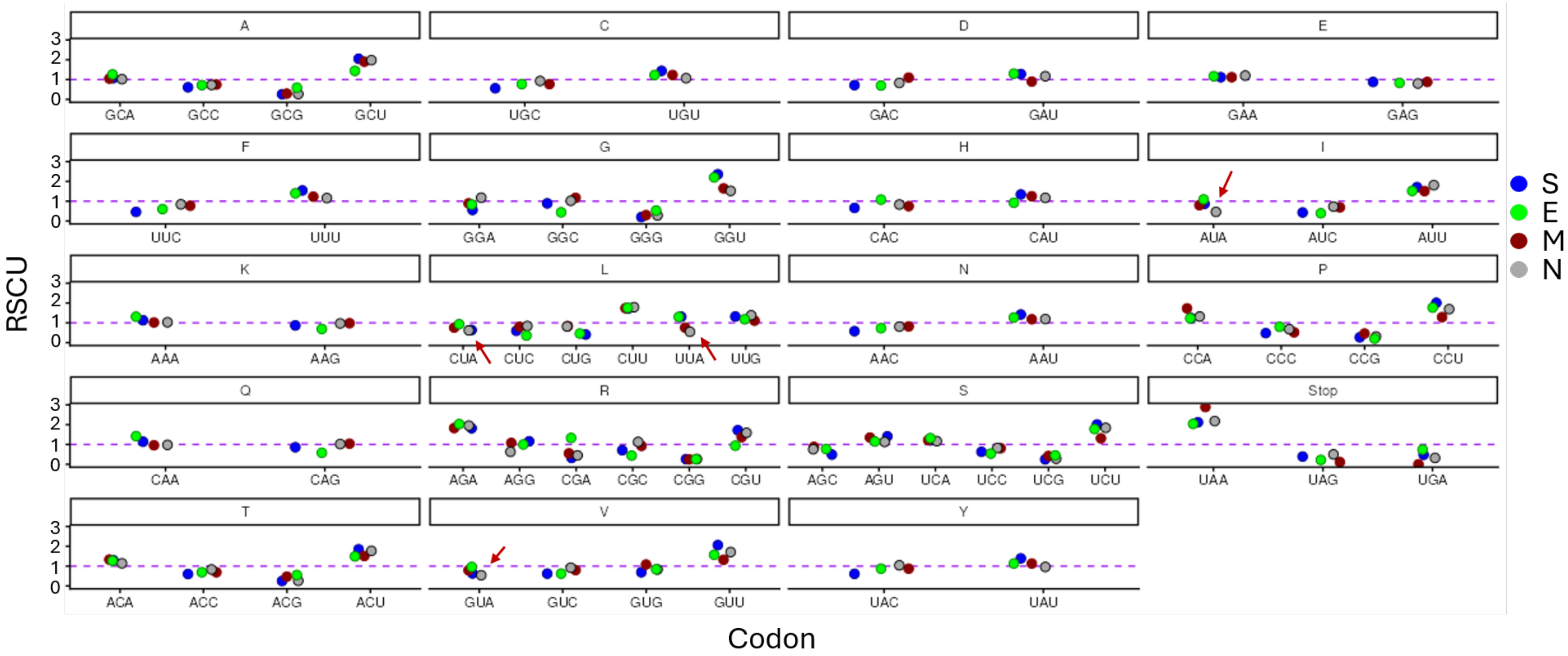


**Supplementary figure 7. Relative synonymous codon usage (RSCU) of CoV structural genes S, E, M and N, grouped by amino acid**. RSCU was calculated on MEGA 11 from codon-based multiple sequence alignments of each structural gene. An RSCU of 1, which indicates random usage, is shown by the dotted line. Red arrows indicate N-U-A codons.

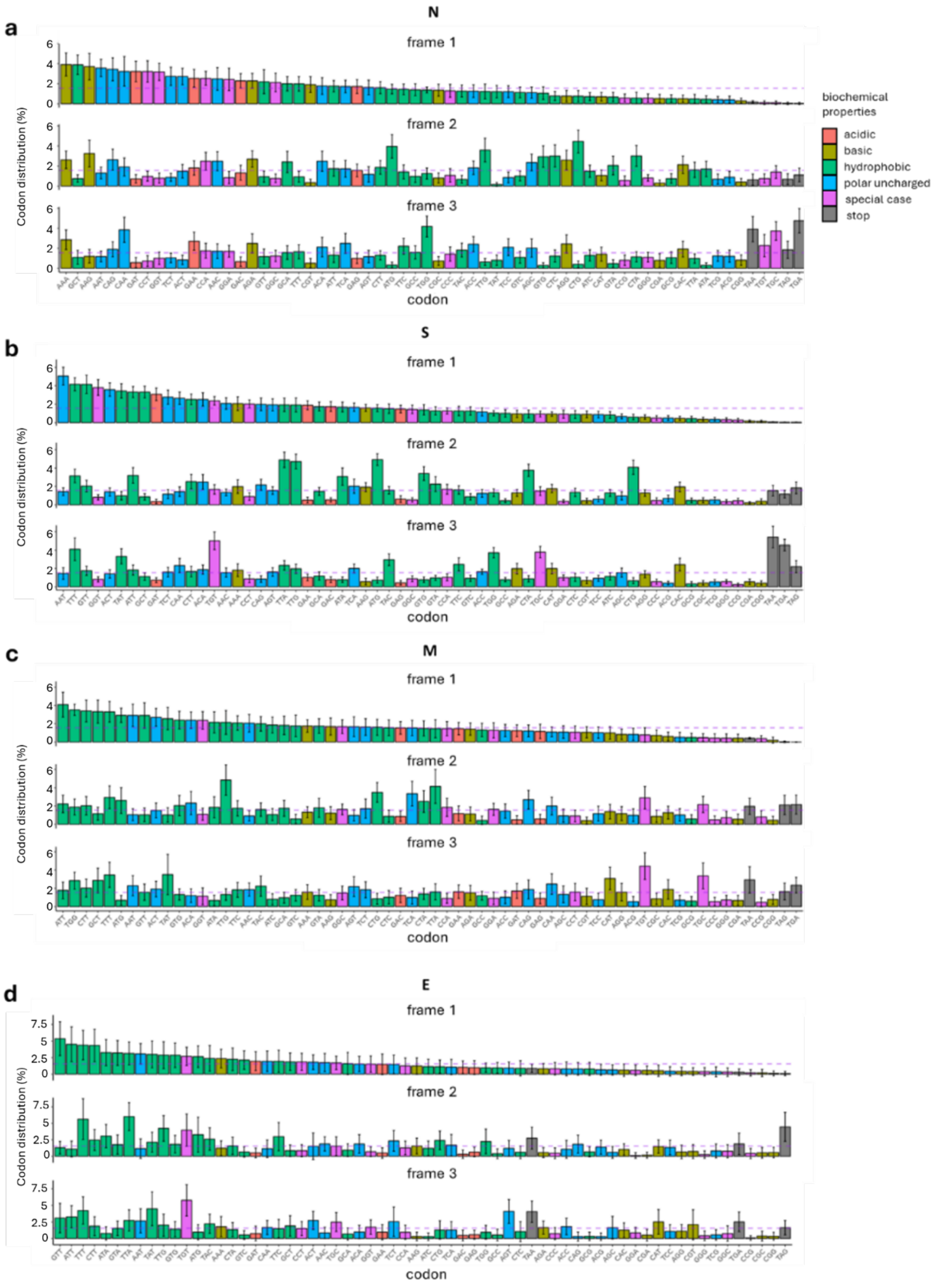


**Supplementary figure 8. Codon distribution of CoV structural genes across plus sense reading frames.** Codon **frequency** distribution for (a) Nucleocapsid, (b) Spike, (c) Membrane and (d) Envelope genes in our dataset, divided by reading frame and colour-coded according to the biochemical properties of the amino acids they encode (acidic = orange; basic = mustard; hydrophobic = green; polar uncharged = blue; special case (C, G, P) = fuchsia; Stop = grey). Error bars represent the standard deviation.

**SUPPLEMENTARY CODE**

**Orfan R script**

```
library(viralORF) library(dplyr) library(magrittr) library(stringr) # ---
Input --- viralORF::find_startstop_ORFs("your_fasta_file.fasta") # find all
ORFs across reading frames of any sequence you want all_ORFs<-
read.csv("viralORF_output.csv") ##dataframe generated using the 'viralORF'
package kozak_reference<-read.csv("kozak_strength_table.csv") #
kozak_reference should include: dinuc, kozak_strength # PRE-PROCESSING
all_ORFs <- all_ORFs %>% mutate(frame = case_when(frame == "plus_0" ~ "1",
frame == "plus_1" ~ "2", frame == "plus_2" ~ "3", TRUE ~ frame)) # Isolate
the +1 and +2 ORFs (candidate AcORFs) plus_1_2_orfs <- all_ORFs %>%
filter(frame %in% c("2", "3")) %>% group_by(seq, end) %>%
arrange(desc(length)) %>% slice(1) %>% ungroup() %>% select(seq, ORF_ID,
start, end, length, nuc_sequence, kozak_dinuc, frame, sense, kozak_strength)
# Isolate the +0 ORFs (reference gene) plus_0_orfs <- all_ORFs %>%
filter(frame == "1") %>% group_by(seq) %>% arrange(desc(length)) %>% slice(1)
%>% ungroup() %>% select( seq, gene_start = start, gene_length = length,
gene_kozak_strength = kozak_strength, gene_seq = nuc_sequence, frame ) #
Combine reference gene and acORFs combined_data <- plus_0_orfs %>%
left_join(plus_1_2_orfs, by = "seq") %>% ungroup() # Get intervening ATG
Kozak strengths get_iATG_kozaks <- function(seq_region, kozak_reference,
gene_strength) { kozak_strengths <- c(if (is.na(gene_strength)) "unknown"
else gene_strength) if (nchar(seq_region) < 6) return(kozak_strengths)
atg_positions <- unlist(gregexpr("ATG", seq_region, fixed = TRUE)) if
(length(atg_positions) == 0 || atg_positions[1] == -1)
return(kozak_strengths) intervening_positions <-
atg_positions[atg_positions > 1] if (length(intervening_positions) > 0) {
for (pos in intervening_positions) { pre <- if (pos >= 3) substr(seq_region,
pos - 3, pos - 3) else "_" post <- if ((pos + 3) <= nchar(seq_region))
substr(seq_region, pos + 3, pos + 3) else "_" dinuc <- paste0(pre, post)
kozak_strength <- kozak_reference$kozak_strength[match(dinuc,
kozak_reference$dinuc)] if (is.na(kozak_strength)) kozak_strength <-
"unknown" kozak_strengths <- c(kozak_strengths, kozak_strength) } }
return(kozak_strengths) } # Convert Kozak qualitative strength to skipping
probability convert_kozak_to_prob <- function(k) { if (is.na(k)) return(1)
k <- tolower(k) if (k == "strong") return(0.15) if (k == "moderate")
return(0.453333333) if (k == "weak") return(0.85) return(1) } #Distance
scoring get_distance_score <- function(distance) { if (is.na(distance))
return(0) if (distance < 100) return(1) if (distance > 300) return(0)
return(0.5) } # Length scoring get_length_score <- function(length) { if
(is.na(length)) return(0) if (length >= 120) return(1) return(0) } # Main
computation combined_data_with_iATGs <- combined_data %>% rowwise() %>%
mutate( unaligned_5prime = if_else( !is.na(gene_start) & !is.na(start) &
(as.numeric(start) > as.numeric(gene_start)), substr(gene_seq, 1,
(as.numeric(start) - as.numeric(gene_start))), "" ), iATG_Vs =
list(get_iATG_kozaks(unaligned_5prime, kozak_reference,
gene_kozak_strength)), unaligned_atg_count = length(iATG_Vs) ) %>% ungroup()
# Expand iATG columns max_iATGs <-
max(lengths(combined_data_with_iATGs$iATG_Vs), na.rm = TRUE) iATG_cols <-
paste0("iATG_V", seq_len(max_iATGs)) for (col in iATG_cols) {
combined_data_with_iATGs[[col]] <- NA_character_ } for (i in
seq_len(nrow(combined_data_with_iATGs))) { vset <-
combined_data_with_iATGs$iATG_Vs[[i]] if (length(vset) > 0) { for (j in
seq_along(vset)) { combined_data_with_iATGs[[iATG_cols[j]]][i] <- vset[j] }
```

```
} } combined_data_with_iATGs$iATG_Vs <- NULL # Compute final numeric scores
and initiation potential combined_data_with_scores <-
combined_data_with_iATGs %>% rowwise() %>% mutate( distance =
if_else(!is.na(start) & !is.na(gene_start), as.numeric(start) -
as.numeric(gene_start), NA_real_), kozak_score =
convert_kozak_to_prob(kozak_strength), initiation_potential = 1 -
kozak_score, distance_score = get_distance_score(distance), length_score =
get_length_score(length), iATG_prob_product = { raw_data <-
c_across(all_of(iATG_cols)) numeric_vals <- vapply(na.omit(raw_data),
function(k) { as.numeric(convert_kozak_to_prob(k)) }, FUN.VALUE =
numeric(1)) if (length(numeric_vals) == 0) 1 else prod(numeric_vals) },
final_score = initiation_potential * distance_score * length_score *
iATG_prob_product ) %>% ungroup() #Save output output_scored_name <-
"gene_scored_ORFs.csv" write.csv(combined_data_with_scores, file =
output_scored_name, row.names = FALSE) cat("Scoring complete.\n") cat("File
saved as:", output_scored_name, "\n")
```

This code was written by FDA and debugged using AI assistance from Google Gemini
